# Detection of low-density *Plasmodium falciparum* infections using amplicon deep sequencing

**DOI:** 10.1101/453472

**Authors:** Angela M. Early, Rachel F. Daniels, Timothy M. Farrell, Jonna Grimsby, Sarah K. Volkman, Dyann F. Wirth, Bronwyn L. MacInnis, Daniel E. Neafsey

## Abstract

**Background:** Deep sequencing of targeted genomic regions is becoming a common tool for understanding the dynamics and complexity of *Plasmodium* infections, but its lower limit of detection is currently unknown. Here, a new amplicon analysis tool, the Parallel Amplicon Sequencing Error Correction (PASEC) pipeline, is used to evaluate the performance of amplicon sequencing on low-density *Plasmodium* DNA samples. Illumina-based sequencing of two *P. falciparum* genomic regions (*CSP* and *SERA2*) was performed on two types of samples: *in vitro* DNA mixtures mimicking low-density infections (1-200 genomes/μl) and extracted blood spots from a combination of symptomatic and asymptomatic individuals (44-653,080 parasites/μl). Three additional analysis tools—DADA2, HaplotypR, and SeekDeep—were applied to both datasets and the precision and sensitivity of each tool were evaluated.

**Results:** Amplicon sequencing can contend with low-density samples, showing reasonable detection accuracy down to a concentration of 5 *Plasmodium* genomes/μl. Due to increased stochasticity and background noise, however, all four tools showed reduced sensitivity and precision on samples with very low parasitemia (<5 copies/μl) or low read count (<100 reads per amplicon). PASEC could distinguish major from minor haplotypes with an accuracy of 90% in samples with at least 30 *Plasmodium* genomes/μl, but only 61% at low *Plasmodium* concentrations (<5 genomes/μl) and 46% at very low read counts (<25 reads per amplicon). The four tools were additionally used on a panel of extracted parasite-positive blood spots from natural malaria infections. While all four identified concordant patterns of complexity of infection (COI) across four sub-Saharan African countries, the COI values obtained for individual samples differed in some cases.

**Conclusions:** Amplicon deep sequencing can be used to determine the complexity and diversity of low-density *Plasmodium* infections. Despite differences in their approach, four state-of-the-art tools resolved known haplotype mixtures with similar sensitivity and precision. Researchers can therefore choose from multiple robust approaches for analyzing amplicon data, however, error filtration approaches should not be uniformly applied across samples of varying parasitemia. Samples with very low parasitemia and very low read count have higher false positive rates and call for read count thresholds that are higher than current recommendations.

## Background

Amplicon deep sequencing is an increasingly utilized genotyping approach that provides a cost-effective strategy to profile the genetic diversity of pathogen infections. Like single nucleotide polymorphism (SNP)-based genotyping methods, both the data-generation and data-analysis steps of amplicon sequencing are highly scalable, allowing for studies of hundreds to thousands of samples. Additionally, amplicons can be designed to cover long genetic segments composed of multiple variants, allowing for the identification of complete DNA sequences (or haplotypes) in a targeted genomic region. When targeting a highly polymorphic genomic region, a single amplicon can distinguish among hundreds of unique haplotypes [1], providing higher resolution than either SNP-based or length-based genotyping approaches. This improves estimates of the number of lineages within polyclonal infections (or complexity of infection; COI) [2–4], permits the discovery of unknown alleles [5–7], and provides increased information for haplotype-based analyses of epistasis and linkage disequilibrium [8].

Amplicon analysis in *Plasmodium* has been adapted to multiple sequencing platforms depending on the desired cost, sample size, and sequence length [3, 9–11]. Because of this high resolution and flexibility, amplicon-based methods have been utilized in a range of applications, including studies of allele-specific vaccine efficacy [1], disease severity [10], clearance rate [12], within-host competition [13], relapse rate [9], drug resistance [5–7], host selection [8], and population structure [8, 14]. Amplicon sequencing has high sensitivity for the detection of minority parasite lineages within an infection, and is of particular interest in longitudinal studies that track intra-host dynamics [3, 4].

When used to detect known single variant markers, amplicon sequences can be analyzed with relatively straightforward approaches. Longer, complex haplotypes, however, require more sophisticated analysis methods. Amplicon sequencing data are known to be subject to PCR and sequencing artifacts, particularly for genomic regions with high A/T-content and high rates of homopolymerism [15, 16]. In addition, library preparation method and primer choice can influence the types and extent of errors [17]. Correctly identifying sequence errors is therefore a challenge when applying amplicon sequencing to *P. falciparum*. Fortunately, several new analytical tools have been developed in recent years to address these challenges [18–21]. Unlike approaches that use reference datasets or cluster sequences with hard percent-identity thresholds, these new methods are more flexible and can distinguish among sequences that differ by only a single nucleotide change [22]. When *Plasmodium* concentrations are reasonably high, these approaches have been demonstrated to be robust. To date, however, none of these methods have been tested on low-density *Plasmodium* samples. It is therefore unclear whether additional considerations are required when interpreting amplicon sequencing data from infections with low parasitemia.

This study assesses amplicon sequencing’s lower limit of detection using four analysis tools, and further evaluates each tool’s accuracy and capacity to recover quantitative information on the relative abundance of different haplotypes within infections. Three of these tools—DADA2 [18], HaplotypR [19], and SeekDeep [20]—were previously published and developed to contend with any *Plasmodium* amplicon. The fourth—the Parallel Amplicon Sequencing Error Correction (PASEC) pipeline—is a distance- and abundance-based error-correction tool that was specifically tailored for use with *CSP* and *SERA2* amplicons [1] and is formally presented here for the first time. Amplicon sequencing of densely polymorphic regions in the *P. falciparum CSP* and *SERA2* genes was applied to two sample collections. The first—a set of *in vitro* human/parasite DNA mixtures that mimic low density parasite infections—was designed to test the limit of detection for amplicon sequencing. The second sample set consisted of DNA extracted from dried blood spots from malaria-infected individuals collected on filter paper in sub-Saharan Africa. This allowed a comparison of analysis approaches using conditions under which samples are typically collected and processed. All four tools detected *P. falciparum* haplotypes with high sensitivity, and additionally were able to discriminate between major and minor haplotypes with reasonable accuracy. Additionally, PASEC was able to identify a *SERA2* indel in patient samples due to its incorporation of prior knowledge on sequence composition. Overall, the results show that low parasitemia does not preclude amplicon analysis of *P. falciparum* samples, although researchers should expect reduced sensitivity and reduced precision with low read-count samples (<100 reads/amplicon) and at parasite densities under 5 genomes/μl.

## Methods

### Sample assembly and composition

#### Mock *Plasmodium*/human DNA mixtures

Mixtures of DNA from cultured *P. falciparum* parasites were combined with human genomic DNA to construct samples that mimic human infections. DNA from up to five culture-adapted parasite lines were combined in various proportions and number (Figure 1; exact sample composition is in Additional File 1, Table S1). Stock mixtures of 200 genomic copies/μl were prepared by real-time PCR quantification of copies/μl in triplicate relative to a plasmid containing a single copy of the quantification target gene [23]. These stock solutions were then diluted to the indicated concentrations in sequencing-grade water and 10 ng commercial human DNA (Promega Corp cat#G3041) was added to all samples. After mixing and dilution, a subset of samples were re-quantified using the same qPCR protocol and reported sample concentrations were adjusted as needed. *Plasmodium-*free negative control samples were also constructed. These contained either 10 ng of human DNA or only water.

**Figure 1.**
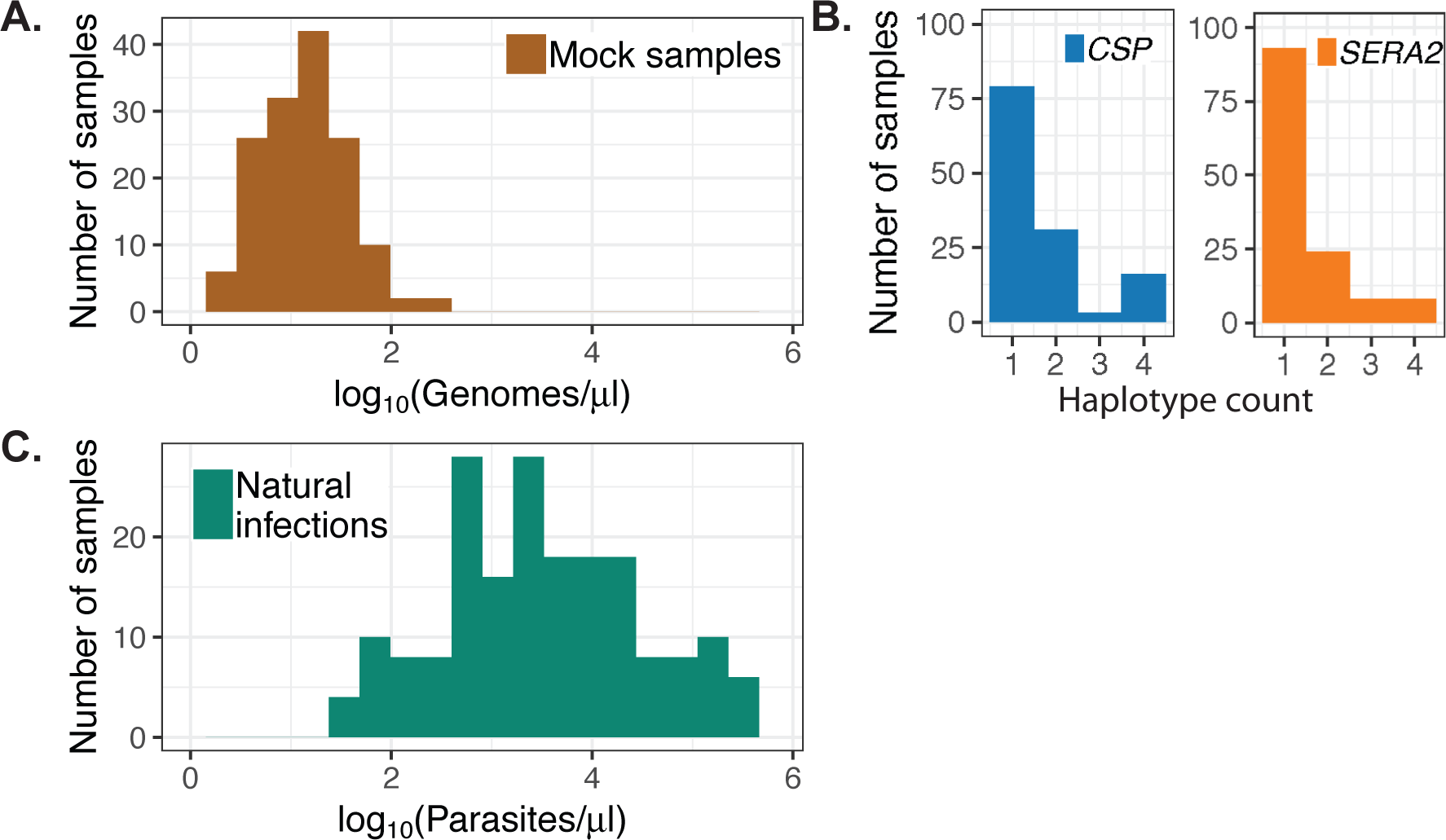
Mock and natural infection sample composition. (A) Mock infection samples were constructed from mixtures of *P. falciparum* and human DNA to mimic the parasite DNA concentrations found in extracted low-density infections. (B) DNA from up to five clonal cultured parasite lines was combined to create each mock sample, leading to within-sample haplotype counts of one to four. (C) Natural infection samples were previously collected and extracted from a combination of symptomatic patients and asymptomatic carriers [1]. Parasite densities were determined by blood smear.

#### Natural infections

Previously extracted DNA from 95 blood spots, obtained from individuals infected with *P. falciparum*, was re-amplified and re-sequenced as part of this study. These samples were acquired from both symptomatic and asymptomatic individuals from four countries in sub-Saharan Africa as part of the RTS,S malaria vaccine Phase 3 trial and had parasite densities that ranged from 44-653,080 parasites/μl as determined by blood smear (Figure 1; [24]). Full details on sampling and extraction, including human subjects approval for use of these samples, are provided in Neafsey *et al.*, 2015 [1]. In brief, samples were collected as blood spots on Whatman FTA cards, shipped to the Broad Institute, and stored in desiccators until processing. DNA was extracted in batches of 95 samples plus one blank control card using the automated Chemagen Chemagic bead-based extraction platform. Total DNA was stored at −80°C until re-amplification and sequencing.

#### Positive control plasmid

A plasmid containing synthetic target amplicon sequences for both *CSP* and *SERA2* was obtained from a commercial vendor (Invitrogen/Thermo Fisher Scientific) and served as a positive control during the PCR amplification step. Outside the primer regions, the plasmid sequence contains nucleotide variants not observed in natural *P. falciparum* isolates so that any instances of contamination can be readily identified. The plasmid map can be found in Additional File 1, Figure S1.

### PCR and sequencing

Two regions from the *CSP* (PF3D7_0304600) and *SERA2* (PF3D7_0207900) genes were PCR amplified as previously described [1]. In brief, 5 μl of DNA were amplified at the targeted regions then indexed in two separate rounds of PCR. The final *CSP* and *SERA2* amplicons cover 288 and 258 nucleotides, respectively (Pf3D7_03_v3:221,352-221,639; Pf3D7_02_v3:320,763-321,020). Both amplicons overlap sequence regions of high nucleotide diversity in sub-Saharan Africa to maximize the number of distinct haplotypes that can be detected across samples from this geographic area.

All DNA samples and negative controls were amplified and sequenced in duplicate. Paired-end 250-bp reads were generated in one MiSeq run conducted on a pool of 384 PCR products. Unless otherwise noted, each PCR/sequencing technical replicate was analyzed as a distinct sample. Before downstream analysis, raw sequencing data were demultiplexed and aligned to amplicon reference sequences to remove all non-*Plasmodium* sequences.

### Sample analysis with PASEC

For each sample, paired-end reads were merged using FLASH [25] and aligned with BWA-MEM v0.7.12-r1039 [26] to the amplicon regions of the *P. falciparum* reference genome assembly (PlasmoDB v.9.0 3D7). Two short homopolymeric tracts in *CSP* were masked from analysis, as such regions are highly error-prone in Illumina sequencing and these specific tracts were not known to harbor natural polymorphisms. Masked coordinates are given in Additional File 3.

Within each sample, haplotypes were filtered according to a set of pre-specified thresholds developed by Neafsey *et al* [1]. Haplotypes were required to (1) cover the entire amplicon region, (2) have no uncalled bases, (3) be supported by at least two sets of merged read pairs (henceforth referred to simply as “reads”), and (4) have an intra-sample frequency ≥ 0.01. To account for potential PCR and sequencing errors, the filtered haplotypes were clustered based on nucleotide distance and read depth. If two haplotypes within the same sample differed by only one nucleotide and had a read coverage ratio ≥8:1, they were merged, maintaining the identity of the more common haplotype. Previous implementations of this pipeline removed all potential chimeric reads and required samples to contain at least 200 reads for one of the two amplicons [1, 8]. In this analysis, these metrics were analyzed, but hard filters were not applied to the samples before downstream analysis.

Full details on the PASEC pipeline, its customizable parameters, and its implementation in this study are found in Additional Files 2 and 3 and at https://github.com/tmfarrell/pasec.

### Sample analysis with DADA2, HaplotypR, and SeekDeep

All samples were independently analyzed using three additional amplicon analysis tools: DADA2 [18], HaplotypR [19], and SeekDeep [20]. Beyond the changes detailed below, input parameters deviated only modestly from the default settings. Parameters and scripts used for executing each pipeline can be found in Additional File 3. While previous implementations of PASEC applied a 200 reads/sample threshold, no read count filters were applied at the sample level in the analysis comparisons.

SeekDeep gives the option of grouping data from technical PCR/sequencing replicates of the same sample and applying clustering and filtering to this grouped data to increase confidence in final calls. The pipeline was therefore run under two conditions: grouping technical replicates (the recommended, default SeekDeep approach; “SeekDeep2x”) and treating each PCR/sequencing replicate independently (“SeekDeep1x”). This permitted more equivalent comparisons among pipelines that do not incorporate replicate information and allowed for a determination of whether a single replicate is sufficient for making accurate haplotype calls.

For HaplotypR, the command-line interface was extended in two ways. First, it was altered to return full haplotype sequences as opposed to only bases at variant positions. Second, the trimming input command was expanded to allow each amplicon to have different lengths. The version of HaplotypR used in this analysis can be found at https://github.com/tmfarrell/HaplotypR. After running the pipeline, the authors’ recommended sample-level filtering was applied to the data. Specifically, each sample was required to have a minimum of 25 reads, and individual haplotypes needed to have a minimum of 3 reads and a within-host frequency of at least 0.1%.

### Comparison of analysis tools

All four tools were assessed for their ability to resolve haplotypes at within-sample frequencies down to 1% using the mock low-parasitemia samples. Two performance metrics were computed by comparing expected vs. observed haplotypes in each sample: sensitivity (proportion of all expected haplotypes that were observed) and precision (proportion of all observed haplotypes that were expected). For sensitivity calculations, only haplotypes present at a concentration of at least 1 copy/μl were considered. For each tool, samples were only included in the performance metric calculation if at least one haplotype was identified. Except for the SeekDeep2x implementation, each PCR/sequencing replicate was analyzed as a distinct sample.

## Results

### Sequencing coverage for low-density mock infections and natural infections from sub-Saharan Africa

In total, 148 DNA mixtures of known haplotypic composition, 190 blood samples from sub-Saharan Africa, 12 positive-control plasmid samples, and 4 negative-control samples without *Plasmodium* DNA were PCR amplified for *CSP* and *SERA2* and sequenced on a single Illumina MiSeq run.

The 148 mock infections were constructed to mimic infections with low parasite density and contained between 1 and 200 *P. falciparum* genomes/μl (Figure 1A). These values roughly correspond to parasite densities of 1 and 200 parasites/μl as mature, multi-nucleated blood-stage parasites are generally absent from sampled peripheral blood. Samples at the lowest end of this distribution (1 genome/μl) should have had, on average, five genomic copies transferred to the initial PCR reaction. After sequencing, 145 samples had full-length read coverage for at least one of the two amplicons. For each amplicon, initial raw coverage across these samples ranged from 0 to 280,876. After implementing the PASEC pipeline, coverage ranged from 0 to 31,787 reads. Coverage was sufficient for both amplicons, although median coverage was higher for *CSP* than for *SERA2* (1872 vs. 909; Figure 2A). All samples with low coverage (<100 reads) had *Plasmodium* DNA concentrations below 21 genomes/μl. Overall, however, coverage and genome copy number were only weakly correlated (Spearman’s ρ = 0.55, *P* = 9.3×10^−14^; Figure 2B), suggesting that stochastic factors influence read counts for low parasitemia samples in general.

**Figure 2.**
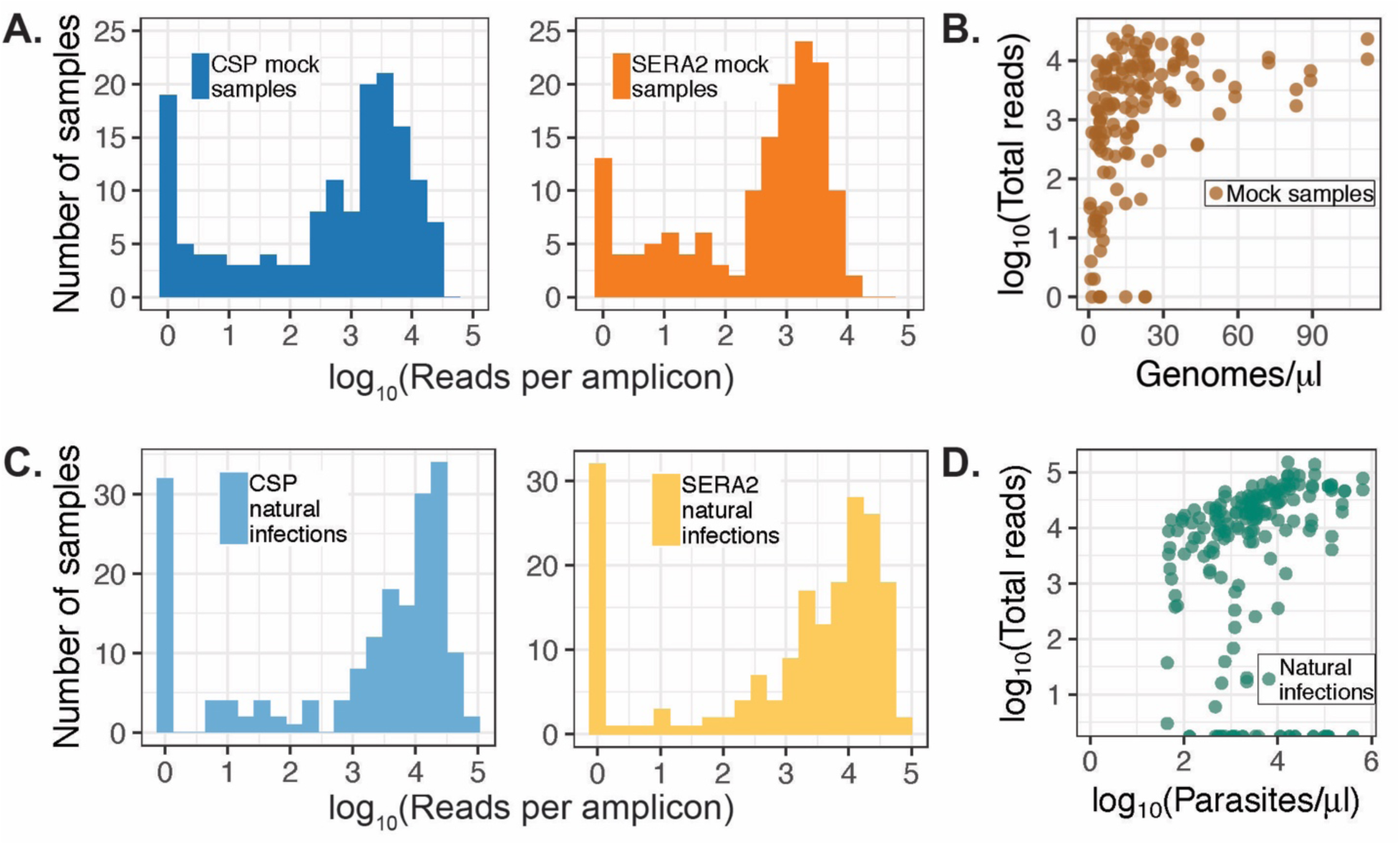
Sequencing coverage of mock and natural infection samples. Overall sequencing coverage was lower for mock infection (A) than natural infection (C) samples (Mann-Whitney U Test, *P* = 1×10^−7^) although natural infections had a higher proportion of samples with no reads. Total read coverage (reads combined from both amplicons) correlated weakly with parasite genome concentration for mock infections (B) and parasitemia for natural infections (D).

Sequence coverage was higher for the samples from natural infections (Figure 2C). These samples were extracted from dried blood spots and had parasite densities that ranged from 44-653,080 parasites/μl as determined by microscopy of blood smears. As with the mock infections, coverage was generally higher for samples with higher parasite loads, but this correlation was low (Spearman’s ρ = 0.31, *P* = 1.1×10^−9^; Figure 2D). While sequencing coverage was higher, overall sequencing success was lower for the natural than for the mock infections (Figure 2C), a likely result of difficulties with extracting high quality DNA from the stored filter paper blood spots. As would be expected under this scenario, failure rate was not evenly distributed across the natural infection samples, suggesting some experienced a higher degree of degradation. Each of the 95 blood samples was PCR amplified and sequenced in duplicate, yielding two *CSP* and two *SERA2* technical replicates per initial blood sample extraction, or 340 total amplicon samples. Of these 340 amplicon samples, 94 (25%) had low read counts (<100 reads). These failures clustered in a small number of blood samples, suggesting that amplification and sequencing success is dependent on sample quality: only 33 (35%) of the blood samples experienced any amplicon failure and 18 samples (19%) received low read counts for all 4 amplicon attempts.

### Absolute haplotype concentration affects the probability of sequencing success

One challenge of amplicon sequencing analysis is to correctly resolve individual haplotypes present within an infection at varying concentrations. Each mock sample contained between one and four unique haplotypes at the *CSP* and *SERA2* amplicons present at concentrations of 1-200 copies/μl (Figure 1B). Overall, there was a high recovery of these expected haplotypes from each of the samples. PASEC correctly identified all haplotypes present at a concentration of 30 copies/μl or higher and 96% of haplotypes with concentrations over 20 copies/μl. Conversely, only 41% of haplotypes with 1-5 copies/μl were recovered (Figure 3A). As discussed in the tool comparison below, this haplotype sensitivity is only slightly influenced by the post-sequencing analysis method and instead is driven by a failure to initially amplify and/or sequence these low frequency haplotypes.

**Figure 3.**
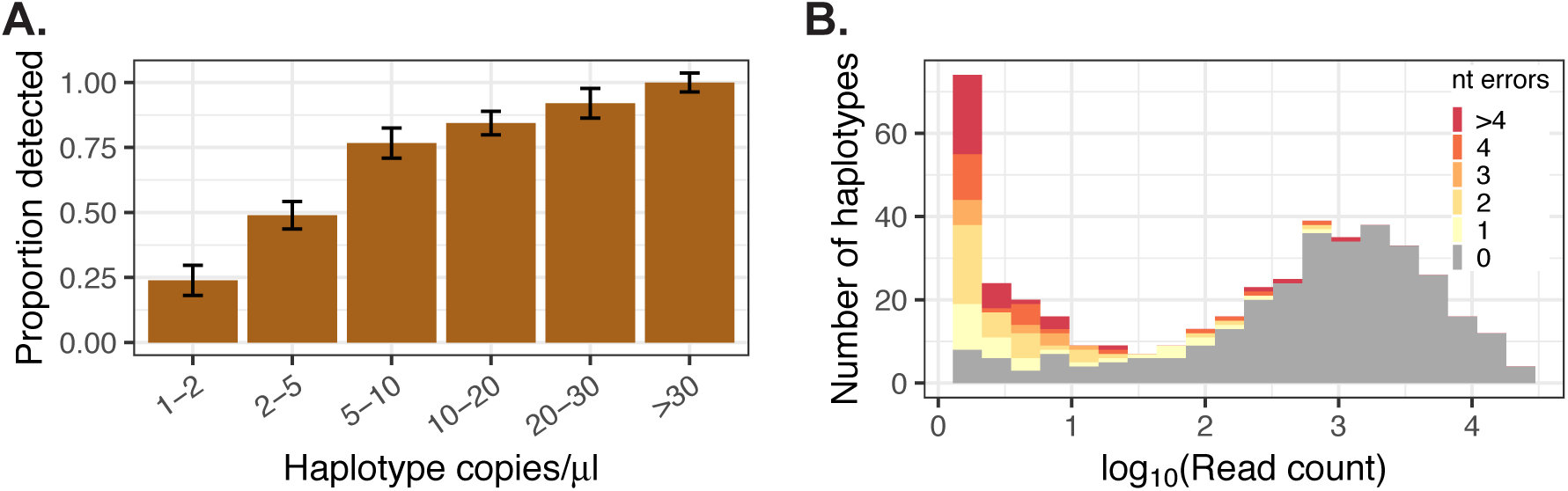
Identification of haplotypes in mock samples. (A) Detection of known haplotypes within the mock samples was dependent on the haplotype concentration (copies/μl) within the sample. Error bars represent the binomial-estimated standard deviation. (B) Across all mock samples, 31% of identified haplotypes were erroneous, but these haplotypes were generally supported by fewer reads than correct haplotypes. The number of nucleotide (nt) errors per haplotype was calculated as the nucleotide distance between an observed haplotype and the closest expected haplotype within the sample.

### Amplicon sequencing retains some information on within-sample haplotype frequencies, even at low concentrations

When performing direct short-read sequencing, relative read depth can be used to infer sample features like genotype ratios or genome copy number variations. During construction of amplicon libraries, however, PCR amplification prior to sequencing introduces stochastic variation in the final read counts. Nevertheless, analysis of the final read ratios in the mock samples shows that some information about the original haplotype ratios can be recovered. For samples with at least 100 reads, the correlation between the haplotypic ratio in the template DNA and final read ratio was moderate (Pearson’s r = 0.82, *P* < 0.001, Additional File 1, Figure S2). As a result, in 73% of samples with at least a 4% margin between the two most prevalent haplotypes, read ratio correctly identified the most prevalent haplotype in the starting DNA mixture. Again, low read count reduced the probability of identifying the correct major haplotype (Figure 4A). Similarly, major haplotype identification was less accurate in samples with very low total *Plasmodium* DNA concentration (<5 genomes/μl; Figure 4B).

**Figure 4.**
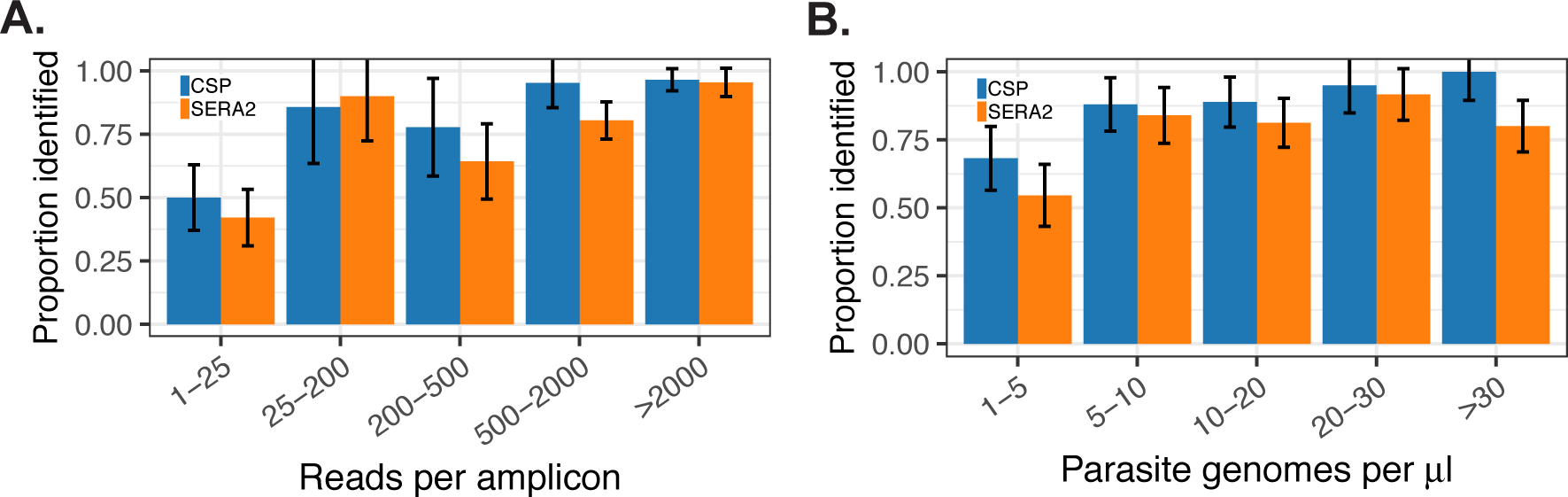
Proportion of mock samples where the major haplotype was correctly identified. Identification of the major haplotype within a sample was less reliable at (A) low read counts and (B) low parasite genome concentrations. Samples were excluded from the analysis if the difference in prevalence between the top two haplotypes was less than 4%. Error bars represent the binomial-estimated standard deviation.

### Erroneous haplotypes have lower read support than correct haplotypes

Read support is a useful indicator of the likelihood that a called haplotype is correct. Haplotypes with single-read support were largely sequencing artifacts, with only 0.030% matching a haplotype sequence known to be present in the sample mixtures. The default PASEC pipeline therefore requires haplotypes to have read support ≥2, a filter that eliminated 89.0% of *CSP* and 85.8% of *SERA2* initially called haplotypes from the dataset.

After filtration with the full PASEC pipeline, some erroneous haplotypes remained, but they continued to show lower read support than true haplotypes (Figure 3B). In the final filtered dataset, 31% of the identified haplotypes were erroneous, although combined these haplotypes only accounted for 0.75% of the total reads. Of note, the same percentage of erroneous reads (0.8%) was previously reported by Hathaway *et al* on a different dataset analyzed with their tool SeekDeep [20]. Reads supporting erroneous haplotypes were more prevalent in samples with low read depth and low parasite concentration (Additional File 1, Figure S3).

In order to decrease the false positive rate, users can increase the read support threshold per haplotype or the minimum read depth per sample. Striving to completely eliminate false positives, however, would decrease sensitivity, especially for low-frequency haplotypes. For instance, 41% of samples contained at least one erroneous haplotype for one of the two amplicons. In 42% of these cases, the most common erroneous haplotype contained higher read support than the least prevalent true haplotype within the sample.

### Frequency and source of haplotype errors in the mock samples

The PASEC pipeline contains customized filtration and error-correction steps to remove erroneous *CSP* and *SERA2* haplotypes. The filtration and error-correction steps in PASEC were designed to address three main sources of erroneous haplotypes: sequencing errors, chimeric reads, and sample contamination. The frequency of these error types and the efficacy of the various PASEC filters are discussed in more detail below.

#### Nucleotide sequence errors

The majority of erroneous haplotypes are expected to result from sequence errors (nucleotide substitutions or indels) that occur during Illumina sequencing or the initial rounds of PCR. The PASEC pipeline accounted for these errors with two approaches: (1) hard masking of error-prone sequence regions and (2) clustering of haplotypes that differed by a single nucleotide and had a read coverage ratio ≥8:1. Hard masking was applied to two homopolymeric regions in *CSP* composed of 9 and 6 poly-Ts. In the raw data, erroneous indels within these two regions were detected in 5.7% and 1.2% of full-length reads. While true indels might occur in these sequences in natural populations, this high artifactual indel rate suggests that inference of variants in these regions would be too unreliable using Illumina sequencing. Compared to masking, the clustering of haplotypes had an even greater impact on reducing nucleotide errors: 57.0% of *CSP* haplotypes and 47.9% of *SERA2* haplotypes were eliminated at this step.

In the final filtered dataset, approximately half of the erroneous haplotypes (51%) differed from a true haplotype by one or two nucleotide changes and were likely the result of Illumina sequencing or PCR errors. As discussed above, these haplotypes were supported by fewer reads than true haplotypes (Figure 3B).

#### Chimeric reads

Chimeric reads are false recombinant haplotypes generated during PCR amplification. While a necessary consideration when performing amplicon sequencing, their overall impact on the mock sample analysis was minimal. Potential chimeras were identified with the isBimera function in DADA2 [18], which identifies all haplotypes that could be constructed from a simple combination of two other haplotypes within the same sample. This analysis flagged 7 *CSP* and 16 *SERA2* samples as containing a total of 36 chimeric haplotypes. Eleven (31%) of the flagged haplotypes were in fact true haplotypes known to be within the given sample. Further analysis showed that 20 of the 25 flagged erroneous haplotypes were only one nucleotide change away from another haplotype in the sample, and the remaining five were related by two nucleotide changes. This suggests that these haplotypes may have resulted from PCR or sequencing error instead of chimeric read formation. Eighteen (78%) of the flagged samples had total read counts under 200, the read threshold previously used with the PASEC pipeline [1]. The increased stochasticity associated with low-read samples may explain why these haplotypes were not merged as part of the PASEC sequencing error filter.

Correctly identifying chimeric reads in natural infections presents an additional challenge, especially in regions of high malaria prevalence where recombination among haplotypes will be higher. Of the 50 most common *CSP* sequences detected in sub-Saharan Africa [8], 38 (76%) were flagged as chimeric combinations by DADA2. Researchers must therefore consider additional factors like population-level haplotype frequency when identifying chimeric reads in natural infections [19, 20].

#### Cross-sample or environmental contamination

A large percentage (49%) of erroneous haplotypes had no evidence of chimerism and were unlikely to have resulted from sequencing errors as they were ≥3 nucleotide changes away from any true haplotype within a given sample. 68% of these haplotypes were present in other samples from the same MiSeq run, suggesting cross-sample or environmental contamination. The remaining haplotypes occurred only once in the whole dataset and may have resulted from environmental contamination. A small amount of cross-sample or environmental contamination was also observed in the negative control samples that contained either water (N=2) or human DNA (N=2). These four *Plasmodium*-free samples contained 5, 7, 16, and 20 reads, respectively. All of these read counts fell well below the 200-read quality threshold previously used with the PASEC pipeline [1].

#### Comparison of PASEC with three state-of-the-art amplicon analysis tools

The performance of PASEC—a pipeline that has been carefully tuned for use with the *CSP* and *SERA2* amplicons in *P. falciparum*—was compared to that of three analysis tools that were developed to be applied to amplicons from any genomic region: DADA2 [18], HaplotypR [19], and SeekDeep [20]. All four of these tools were designed to detect low-frequency haplotypes and differentiate unique haplotypes with single-nucleotide resolution. There are, however, differences in the analytical approaches. For instance, during error filtration PASEC and HaplotypR rely mainly on variant frequency and read depth, while SeekDeep incorporates k-mer frequencies and base quality scores and DADA2 further models sequencer-specific error likelihoods. SeekDeep additionally allows users to incorporate replicate PCR and sequencing runs into the analysis. This approach provides higher confidence for differentiating between sequencing errors and true haplotypes that differ at only a single nucleotide. As all haplotypes used in the mock samples differed by more than one nucleotide, however, this SeekDeep feature was not evaluated in the trial.

While all these tools have undergone rigorous testing, no previous study has focused on their performance under extremely low parasite densities. Here, each tool was applied to the mock samples and it was evaluated on (1) the proportion of all expected haplotypes that were observed (sensitivity) and (2) the proportion of observed haplotypes that were expected (precision).

#### Sensitivity and precision

Overall, the four tools performed comparably on the mock sample panel, although they showed more variability in precision than in sensitivity (Figure 5). This shows that what differs most between pipelines is their ability to filter out erroneous haplotypes, not identify correct haplotypes. For instance, while the sensitivity of SeekDeep1x—the SeekDeep implementation using only one technical replicate— was comparable to the other four pipelines, its precision was substantially lower, driven by the identification of a high number of erroneous haplotypes. The use of replicate samples in SeekDeep2x greatly decreased the tool’s false positive rate, increasing precision with a small cost in sensitivity.

**Figure 5.**
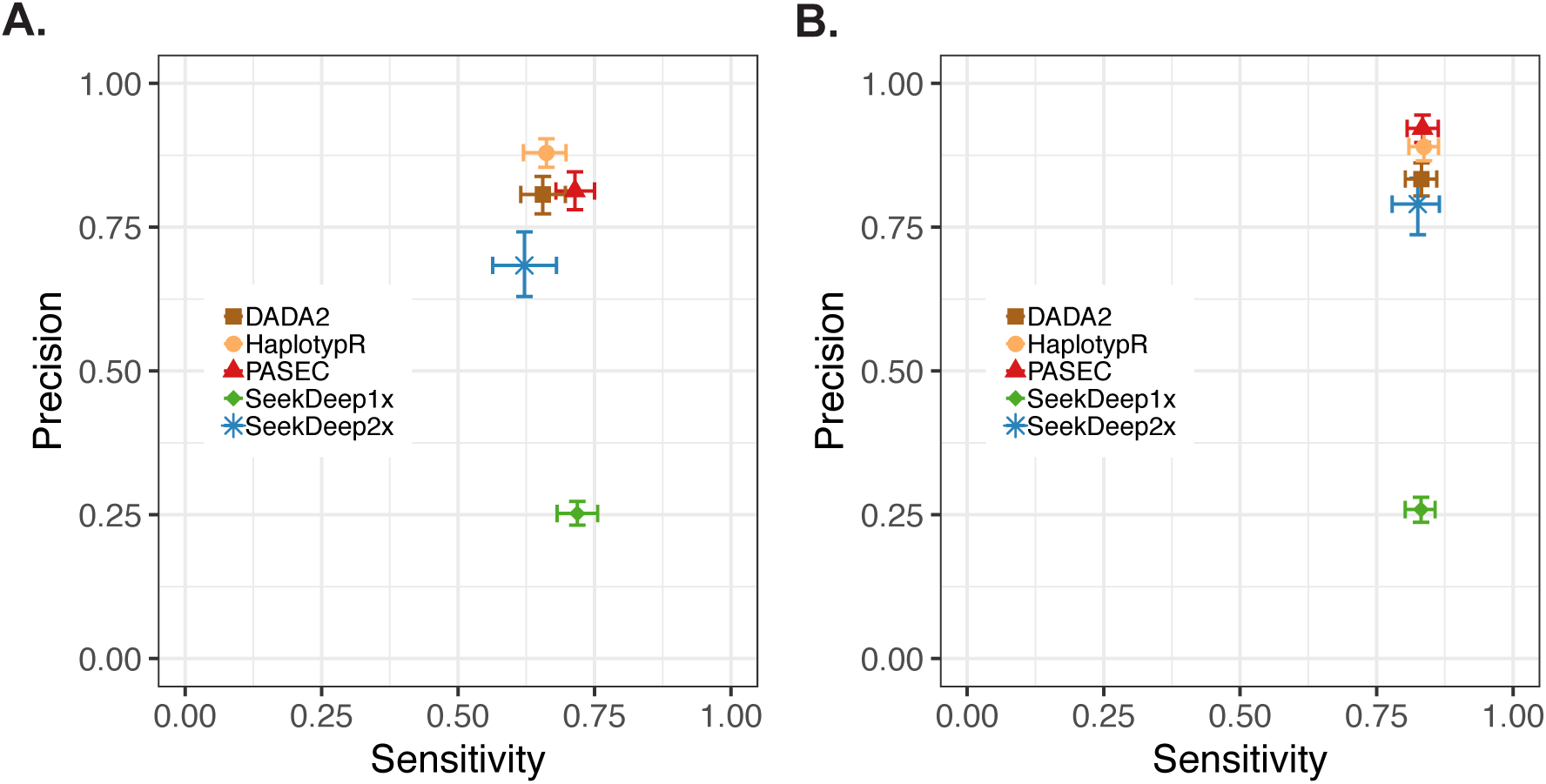
Sensitivity and precision of five analysis pipelines for the detection of haplotypes in mock samples. (A) Analysis approaches vary more in precision than in sensitivity. (B) Performance of all pipelines improves when considering only samples that had at least 100 reads for an individual amplicon. Data shown include results from both the *CSP* and *SERA2* amplicons. 95% confidence intervals were estimated with 1000 bootstrapped data set replicates.

Each tool’s performance varied to some extent across amplicons. This variation was not consistent across pipelines, and as a result, the pipelines’ rank order for precision and sensitivity was different for *CSP* and *SERA2* (Table 1; Additional File 1, Figure S4).

**Table 1.** Sensitivity and precision of each pipeline (Mean [95% CI])

|  |  |  | DADA2 | HaplotypR | PASEC | SeekDeep1x | SeekDeep2x |
| --- | --- | --- | --- | --- | --- | --- | --- |
| All Samples | Sensitivity | All | 0.66 [0.62, 0.70] | 0.66 [0.62, 0.70] | 0.71 [0.68, 0.75] | 0.72 [0.68, 0.76] | 0.62 [0.56, 0.68] |
|  |  | CSP | 0.66 [0.61, 0.71] | 0.64 [0.59, 0.70] | 0.70 [0.64, 0.75] | 0.70 [0.65, 0.75] | 0.61 [0.53, 0.69] |
|  |  | SERA2 | 0.65 [0.59, 0.70] | 0.68 [0.62, 0.74] | 0.73 [0.68, 0.78] | 0.73 [0.68, 0.79] | 0.63 [0.55, 0.71] |
|  | Precision | All | 0.81 [0.77, 0.84] | 0.88 [0.85, 0.90] | 0.81 [0.78, 0.85] | 0.25 [0.23, 0.27] | 0.68 [0.63, 0.74] |
|  |  | CSP | 0.72 [0.67, 0.77] | 0.94 [0.91, 0.97] | 0.86 [0.81, 0.89] | 0.26 [0.23, 0.28] | 0.77 [0.69, 0.84] |
|  |  | SERA2 | 0.91 [0.87, 0.94] | 0.82 [0.78, 0.86] | 0.77 [0.72, 0.82] | 0.25 [0.22, 0.28] | 0.61 [0.53, 0.68] |
| Samples with $\geq 100$ reads | Sensitivity | All | 0.83 [0.80, 0.86] | 0.84 [0.81, 0.86] | 0.83 [0.81, 0.86] | 0.83 [0.80, 0.86] | 0.78 [0.78, 0.87] |
|  |  | CSP | 0.82 [0.78, 0.86] | 0.82 [0.77, 0.86] | 0.82 [0.78, 0.86] | 0.82 [0.78, 0.86] | 0.84 [0.78, 0.89] |
|  |  | SERA2 | 0.85 [0.80, 0.89] | 0.86 [0.81, 0.90] | 0.85 [0.80, 0.89] | 0.85 [0.81, 0.89] | 0.82 [0.75, 0.88] |
|  | Precision | All | 0.83 [0.80, 0.86] | 0.89 [0.87, 0.92] | 0.92 [0.90, 0.94] | 0.26 [0.24, 0.28] | 0.79 [0.74, 0.84] |
|  |  | CSP | 0.75 [0.70, 0.79] | 0.94 [0.91, 0.96] | 0.95 [0.92, 0.97] | 0.27 [0.24, 0.30] | 0.88 [0.83, 0.93] |
|  |  | SERA2 | 0.92 [0.88, 0.95] | 0.84 [0.80, 0.88] | 0.90 [0.86, 0.93] | 0.25 [0.22, 0.28] | 0.71 [0.63, 0.78] |

#### Effect of sample read depth and genome copy number

All five pipelines showed reduced performance at very low read depths (<25 reads/sample) and low parasite concentrations (<5 genomes/μl; Additional File 1, Figure S5). In particular, SeekDeep2x performed best on samples with at least 100 reads (Figure 5B). Parasite genome copy number also affected the tools’ success at resolving at least one haplotype within a sample. Overall, the pipelines reported haplotypes within 78% (HaplotypR), 81% (DADA2), 84% (SeekDeep2x), 89% (PASEC), and 96% (SeekDeep1x) of the samples (Additional File 1, Figure S6A). The majority of the samples returning no data contained *Plasmodium* DNA concentrations under 5 genomes/μl (Additional File 1, Figure S6B).

#### Determination of major haplotype frequency

As reported above, PASEC correctly identified the expected major haplotype in 73% of the mock samples. Misidentification of the expected haplotype could result from errors in the pipeline or stochasticity during sample construction, PCR amplification and sequencing. Strongly suggesting that stochasticity in sample processing and sequencing plays a role, the frequency estimate for each sample’s major haplotype was highly correlated between tools (Pearson’s r for all pairs > 0.85, *P* < 0.001; Additional File 1, Figure S7A). The correlation between tools was even higher when limiting the analysis to samples with at least 100 reads (Pearson’s r for all pairs > 0.97, *P* < 0.001; Additional File 1, Figure S7B). All tools therefore arrive at comparable frequency estimates based on the number of reads produced per haplotype.

### Analysis of natural infection samples from Sub-Saharan Africa with the four tools

All five pipelines were then applied to newly generated amplicon data from 95 previously extracted parasite positive blood spots from four countries in sub-Saharan Africa (Figure 1C) [1]. These biological samples were PCR amplified and sequenced in duplicate, yielding 190 independently sequenced samples for each of the two amplicons. With the exception of SeekDeep2x, the technical replicates were again treated as separate samples in the analysis step. All tools were run with the same parameters used for the mock samples.

The tools differed in the total number of unique haplotypes identified across the samples, with estimates ranging from 48 to 336 for *CSP* and 38 to 412 for *SERA2* (Additional File 1, Figure S8). For both amplicons, SeekDeep1x and DADA2 identified substantially more haplotypes than the other approaches, although a large percentage of these haplotypes were found at within-sample frequencies under 1%, raising the possibility that they were artifacts. Only PASEC identified a three nucleotide indel in *SERA2* that was found on seven different haplotypic backgrounds. This was because the PASEC hard filters permitted this indel to remain based on its prior observation in African parasites [1].

Consistent with expectations for sub-Saharan Africa, the majority of the natural infection samples contained multiple *P. falciparum* parasite haplotypes. COI was estimated for each sample as the maximum number of unique haplotypes identified at either of the two amplicons. With the exception of SeekDeep1x, all four tools produced similar trends of mean COI per country (Figure 6). This is in keeping with the observation that SeekDeep showed lower precision on the mock samples than the other tools when run with single replicates (Figure 5).

**Figure 6.**
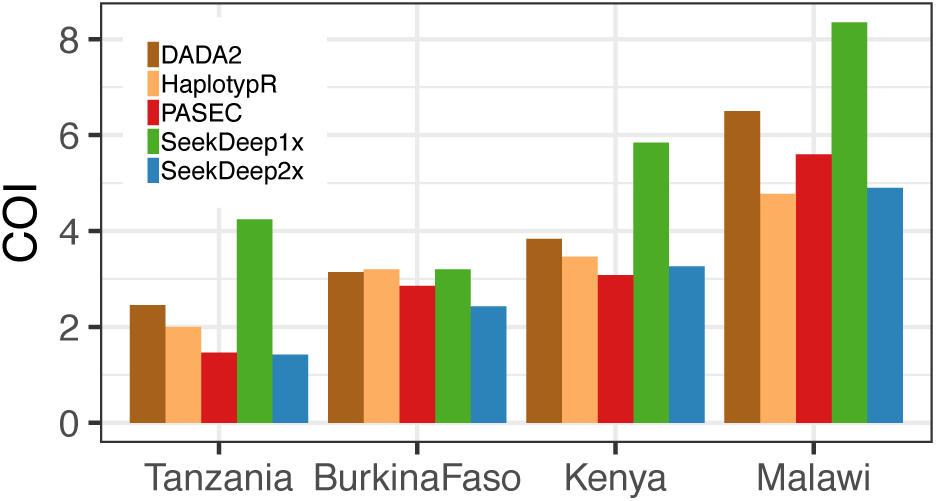
Mean COI estimates for four sub-Saharan African study sites made by the five analysis pipelines. COI was defined as the maximum number of haplotypes retrieved for the sample from either of the two amplicons.

## Discussion

Amplicon sequencing of complex haplotypic regions is a powerful tool being applied to an increasing range of questions in malaria research. This highly scalable approach accurately estimates COI, identifies distinct haplotypes within polyclonal infections, and permits temporal tracking of distinct clones. Previous applications and evaluations of amplicon sequencing have focused on moderate to high density infections. Here, the performance of amplicon sequencing was assessed for the first time under a scenario of extremely low parasite densities (1-200 genomes/μl), which mimicked samples that could be obtained from asymptomatic carriers. The results show that amplicon sequencing remains a viable approach under such challenging scenarios, as it was able to detect 77% of individual haplotypes present at concentrations of 5-10 genomic copies/μl. The ability of Illumina-based amplicon sequencing to reliably detect *Plasmodium* DNA at these extremely low concentrations shows that it has a limit of detection on par with standard nested PCR [27] and qPCR [28] methods.

While amplicon sequencing is successful at low parasite densities, analysis of such samples presents unique challenges, particularly at densities below 5 genomes/μl. At these ultra-low concentrations, overall sample-level error rates are higher and quantification of haplotype ratios is less accurate, regardless of the applied analysis tool. Researchers should therefore take steps to lower false positive rates in this challenging class of samples. Since erroneous haplotypes are generally supported by fewer reads (Figure 3B) and samples with lower read counts have a higher proportion of false haplotypes (Additional File 1, Figure S3), it should be standard practice to raise read thresholds when analyzing low parasitemia or low coverage samples.

PASEC’s high performance was the result of hand-tuning for use with the amplicons *CSP* and *SERA2*. This included the hard masking of difficult-to-sequence homopolymer runs in the *CSP* amplicon and the *a priori* identification of indels in *SERA2*. As a result of this customization, it was the only tool to identify a naturally occurring three nucleotide deletion in *SERA2* that is present in Africa. Importantly, however, this study shows that three other tools—DADA2, HaplotypR, and SeekDeep—also provide robust results when prior knowledge of the error profile of an individual amplicon is unavailable and rapid, parallelized analysis is not needed.

While the overall approach is successful, room for methodological development remains. Improvements in sensitivity will require changes upstream of the analysis stage as the inability to detect a haplotype generally resulted from a failure to capture it at the amplification or sequencing stage. This led to roughly equivalent sensitivities for the four evaluated tools. Precision did vary among tools, reflecting their different approaches towards error correction. As the rank order of the tools’ precision differed between the amplicons, however, the relative success of these different approaches seems dependent on genetic context. With PASEC and SeekDeep, users can easily increase precision by implementing a simple 100 read threshold at the sample level (Table 1). Additional increases in precision will require further development, and work in this area is ongoing [29, 30].

## Conclusion

Amplicon sequencing is a versatile approach for exploring a range of intra-host questions in malaria research. Cost-effective and scalable for use with thousands—or tens of thousands—of polyclonal samples in high-throughput settings, its use will likely increase in the coming years. As shown here, amplicon sequencing can be applied to samples with both low and high parasite densities, although the consistent detection of parasite clones with very low prevalence (<5 genomes/μl) is challenging. Even at low densities, amplicon sequencing retained some information on haplotype ratio, allowing PASEC to distinguish major and minor clones correctly in 73% of the infections. Erroneous haplotypes were generally supported by fewer reads, and samples with lower read counts had a higher proportion of false haplotypes. When used under their recommended conditions, three other versatile analysis tools (DADA2, HaplotypR, and SeekDeep) showed similar performance compared to PASEC. Overall, all tools performed well, and so final choice of analysis method will depend largely on study design (*e.g.*, the inclusion of technical PCR/sequencing replicates), the read coverage of the samples, and expectations regarding the targeted *Plasmodium* genotypes (*e.g.*, the potential presence of indels or the need to differentiate between low frequency haplotypes with a single SNP difference). Regardless of the tool used, however, it should be standard practice to raise read thresholds when analyzing samples with low parasitemia (<5 parasites/μl) or low coverage (<100 reads).

## Acknowledgements

We thank Anita Lerch for thoughtful comments on the manuscript, Katelyn Durfee for help preparing the mock samples, and members of the Neafsey and Wirth labs for helpful discussions throughout the course of the project.

## References

1. Neafsey DE, Juraska M, Bedford T, Benkeser D, Valim C, Griggs A, et al. Genetic Diversity and Protective Efficacy of the RTS,S/AS01 Malaria Vaccine. N Engl J Med. 2015;373:2025–37. doi:10.1056/NEJMoa1505819.

2. Zhong D, Koepfli C, Cui L, Yan G. Molecular approaches to determine the multiplicity of Plasmodium infections. Malar J. 2018;17:172. doi:10.1186/s12936-018-2322-5.

3. Juliano JJ, Porter K, Mwapasa V, Sem R, Rogers WO, Ariey F, et al. Exposing malaria in-host diversity and estimating population diversity by capture-recapture using massively parallel pyrosequencing. Proc Natl Acad Sci. 2010;107:20138–43. doi:10.1073/pnas.1007068107.

4. Lerch A, Koepfli C, Hofmann NE, Kattenberg JH, Rosanas-Urgell A, Betuela I, et al. Longitudinal tracking of Plasmodium falciparum clones in complex infections by amplicon deep sequencing. bioRxiv. 2018;:306860. doi:10.1101/306860.

5. Ngondi JM, Ishengoma DS, Doctor SM, Thwai KL, Keeler C, Mkude S, et al. Surveillance for sulfadoxine-pyrimethamine resistant malaria parasites in the Lake and Southern Zones, Tanzania, using pooling and next-generation sequencing. Malar J. 2017;16:236. doi:10.1186/s12936-017-1886-9.

6. Rao PN, Uplekar S, Kayal S, Mallick PK, Bandyopadhyay N, Kale S, et al. A Method for Amplicon Deep Sequencing of Drug Resistance Genes in Plasmodium falciparum Clinical Isolates from India. J Clin Microbiol. 2016;54:1500–11. doi:10.1128/JCM.00235-16.

7. Talundzic E, Ndiaye YD, Deme AB, Olsen C, Patel DS, Biliya S, et al. Molecular Epidemiology of Plasmodium falciparum kelch13 Mutations in Senegal Determined by Using Targeted Amplicon Deep Sequencing. Antimicrob Agents Chemother. 2017;61:AAC.02116-16. doi:10.1128/AAC.02116-16.

8. Early AM, Lievens M, MacInnis BL, Ockenhouse CF, Volkman SK, Adjei S, et al. Host-mediated selection impacts the diversity of Plasmodium falciparum antigens within infections. Nat Commun. 2018;9:1381. doi:10.1038/s41467-018-03807-7.

9. Lin JT, Hathaway NJ, Saunders DL, Lon C, Balasubramanian S, Kharabora O, et al. Using Amplicon Deep Sequencing to Detect Genetic Signatures of *Plasmodium vivax* Relapse. J Infect Dis. 2015;212:999–1008. doi:10.1093/infdis/jiv142.

10. Patel JC, Hathaway NJ, Parobek CM, Thwai KL, Madanitsa M, Khairallah C, et al. Increased risk of low birth weight in women with placental malaria associated with P. falciparum VAR2CSA clade. Sci Rep. 2017;7:7768. doi:10.1038/s41598-017-04737-y.

11. Dara A, Travassos MA, Adams M, Schaffer DeRoo S, Drábek EF, Agrawal S, et al. A new method for sequencing the hypervariable Plasmodium falciparum gene var2csa from clinical samples. Malar J. 2017;16:343. doi:10.1186/s12936-017-1976-8.

12. Mideo N, Bailey JA, Hathaway NJ, Ngasala B, Saunders DL, Lon C, et al. A deep sequencing tool for partitioning clearance rates following antimalarial treatment in polyclonal infections. Evol Med Public Heal. 2016;2016:21–36. doi:10.1093/emph/eov036.

13. Nair S, Li X, Arya GA, McDew-White M, Ferrari M, Nosten F, et al. Do fitness costs explain the rapid spread of *kelch13*-C580Y substitutions conferring artemisinin resistance? Antimicrob Agents Chemother. 2018;:AAC.00605-18. doi:10.1128/AAC.00605-18.

14. Miller RH, Hathaway NJ, Kharabora O, Mwandagalirwa K, Tshefu A, Meshnick SR, et al. A deep sequencing approach to estimate Plasmodium falciparum complexity of infection (COI) and explore apical membrane antigen 1 diversity. Malar J. 2017;16:490. doi:10.1186/s12936-017-2137-9.

15. Ross MG, Russ C, Costello M, Hollinger A, Lennon NJ, Hegarty R, et al. Characterizing and measuring bias in sequence data. Genome Biol. 2013;14:R51. doi:10.1186/gb-2013-14-5-r51.

16. Schirmer M, D’Amore R, Ijaz UZ, Hall N, Quince C. Illumina error profiles: resolving fine-scale variation in metagenomic sequencing data. BMC Bioinformatics. 2016;17:125. doi:10.1186/s12859-016-0976-y.

17. Schirmer M, Ijaz UZ, D’Amore R, Hall N, Sloan WT, Quince C. Insight into biases and sequencing errors for amplicon sequencing with the Illumina MiSeq platform. Nucleic Acids Res. 2015;43:e37–e37. doi:10.1093/nar/gku1341.

18. Callahan BJ, McMurdie PJ, Rosen MJ, Han AW, Johnson AJA, Holmes SP. DADA2: High-resolution sample inference from Illumina amplicon data. Nat Methods. 2016;13:581–3. doi:10.1038/nmeth.3869.

19. Lerch A, Koepfli C, Hofmann NE, Messerli C, Wilcox S, Kattenberg JH, et al. Development of amplicon deep sequencing markers and data analysis pipeline for genotyping multi-clonal malaria infections. BMC Genomics. 2017;18:864. doi:10.1186/s12864-017-4260-y.

20. Hathaway NJ, Parobek CM, Juliano JJ, Bailey JA. SeekDeep: single-base resolution de novo clustering for amplicon deep sequencing. Nucleic Acids Res. 2018;46:e21–e21. doi:10.1093/nar/gkx1201.

21. Rask TS, Petersen B, Chen DS, Day KP, Pedersen AG. Using expected sequence features to improve basecalling accuracy of amplicon pyrosequencing data. BMC Bioinformatics. 2016;17:176. doi:10.1186/s12859-016-1032-7.

22. Callahan BJ, McMurdie PJ, Holmes SP. Exact sequence variants should replace operational taxonomic units in marker-gene data analysis. ISME J. 2017;11:2639–43. doi:10.1038/ismej.2017.119.

23. Daniels R, Volkman SK, Milner DA, Mahesh N, Neafsey DE, Park DJ, et al. A general SNP-based molecular barcode for Plasmodium falciparum identification and tracking. Malar J. 2008;7:223. doi:10.1186/1475-2875-7-223.

24. RTS,S Clinical Trials Partnership, Agnandji ST, Lell B, Fernandes JF, Abossolo BP, Methogo BGNO, et al. A Phase 3 Trial of RTS,S/AS01 Malaria Vaccine in African Infants. N Engl J Med. 2012;367:2284–95. doi:10.1056/NEJMoa1208394.

25. Magoc T, Salzberg SL. FLASH: fast length adjustment of short reads to improve genome assemblies. Bioinformatics. 2011;27:2957–63. doi:10.1093/bioinformatics/btr507.

26. Li H, Durbin R. Fast and accurate short read alignment with Burrows-Wheeler transform. Bioinformatics. 2009;25:1754–60. doi:10.1093/bioinformatics/btp324.

27. Snounou G, Viriyakosol S, Xin Ping Zhu, Jarra W, Pinheiro L, do Rosario VE, et al. High sensitivity of detection of human malaria parasites by the use of nested polymerase chain reaction. Mol Biochem Parasitol. 1993;61:315–20. doi:10.1016/0166-6851(93)90077-B.

28. Taylor SM, Juliano JJ, Trottman PA, Griffin JB, Landis SH, Kitsa P, et al. High-Throughput Pooling and Real-Time PCR-Based Strategy for Malaria Detection. J Clin Microbiol. 2010;48:512–9. doi:10.1128/JCM.01800-09.

29. Larsson AJM, Stanley G, Sinha R, Weissman IL, Sandberg R. Computational correction of index switching in multiplexed sequencing libraries. Nat Methods. 2018;15:305–7. doi:10.1038/nmeth.4666.

30. Davis NM, Proctor D, Holmes SP, Relman DA, Callahan BJ. Simple statistical identification and removal of contaminant sequences in marker-gene and metagenomics data. bioRxiv. 2018;:221499. doi:10.1101/221499.

