## Supplemental File 1 for "Detection of low-density *Plasmodium falciparum* infections using amplicon deep sequencing"

**Additional File 1: Supplementary Tables and Figures**

**Early *et al*., Amplicon deep sequencing of low-density *Plasmodium falciparum* infections: an evaluation of analysis approaches**

**Table S1: Strain composition of mock *Plasmodium*/human DNA mixtures**

| Total genome copies/ul | Strain Proportion | | | | | |
|---|---|---|---|---|---|---|
| | 3d7 | Mcamp | Th029.09 | Th002.09 | Th135.09 | Dd2 |
| 1.34 | 0 | 1 | 0 | 0 | 0 | 0 |
| 2.18 | 0 | 1 | 0 | 0 | 0 | 0 |
| 2.21 | 0 | 0 | 0 | 0.04 | 0.96 | 0 |
| 2.65 | 0.89 | 0.11 | 0 | 0 | 0 | 0 |
| 2.98 | 0 | 0 | 0 | 0.52 | 0.48 | 0 |
| 3.02 | 0 | 1 | 0 | 0 | 0 | 0 |
| 3.47 | 0 | 0 | 0 | 0.04 | 0.96 | 0 |
| 3.67 | 0.96 | 0.04 | 0 | 0 | 0 | 0 |
| 4.16 | 0.89 | 0.11 | 0 | 0 | 0 | 0 |
| 4.29 | 0.99 | 0.01 | 0 | 0 | 0 | 0 |
| 4.31 | 0 | 0 | 0 | 0 | 1 | 0 |
| 4.50 | 0.99 | 0.01 | 0 | 0 | 0 | 0 |
| 4.67 | 0 | 0 | 0 | 0.52 | 0.48 | 0 |
| 4.7 | 0 | 1 | 0 | 0 | 0 | 0 |
| 4.70 | 1 | 0 | 0 | 0 | 0 | 0 |
| 5.04 | 0 | 0 | 0 | 0.04 | 0.96 | 0 |
| 5.77 | 0.96 | 0.04 | 0 | 0 | 0 | 0 |
| 6.03 | 0.89 | 0.11 | 0 | 0 | 0 | 0 |
| 6.74 | 0.99 | 0.01 | 0 | 0 | 0 | 0 |
| 6.78 | 0 | 0 | 0 | 0 | 1 | 0 |
| 6.79 | 0 | 0 | 0 | 0.52 | 0.48 | 0 |
| 7.06 | 0.99 | 0.01 | 0 | 0 | 0 | 0 |
| 7.39 | 1 | 0 | 0 | 0 | 0 | 0 |
| 8.17 | 0 | 0 | 0 | 0.04 | 0.96 | 0 |
| 8.38 | 0.96 | 0.04 | 0 | 0 | 0 | 0 |
| 9.79 | 0.99 | 0.01 | 0 | 0 | 0 | 0 |
| 9.79 | 0.89 | 0.11 | 0 | 0 | 0 | 0 |
| 9.84 | 0 | 0 | 0 | 0 | 1 | 0 |
| 10.03 | 0 | 0.19 | 0 | 0 | 0.81 | 0 |
| 10.26 | 0.99 | 0.01 | 0 | 0 | 0 | 0 |
| 10.73 | 1 | 0 | 0 | 0 | 0 | 0 |
| 11.01 | 0 | 0 | 0 | 0.52 | 0.48 | 0 |
| 11.30 | 0 | 0 | 0 | 0.04 | 0.96 | 0 |
| 13.55 | 0.89 | 0.11 | 0 | 0 | 0 | 0 |
| 13.60 | 0.96 | 0.04 | 0 | 0 | 0 | 0 |
| 14.75 | 0.33 | 0.04 | 0 | 0.33 | 0.30 | 0 |
| 14.88 | 0.32 | 0.04 | 0.04 | 0.32 | 0.29 | 0 |

| | | | | | | |
|---|---|---|---|---|---|---|
| 15.23 | 0 | 0 | 0 | 0.52 | 0.48 | 0 |
| 15.88 | 0.99 | 0.01 | 0 | 0 | 0 | 0 |
| 15.96 | 0 | 0 | 0 | 0 | 1 | 0 |
| 16.64 | 0.99 | 0.01 | 0 | 0 | 0 | 0 |
| 17.4 | 0 | 0 | 0 | 1 | 0 | 0 |
| 17.4 | 1 | 0 | 0 | 0 | 0 | 0 |
| 17.61 | 0 | 0 | 0 | 0.04 | 0.96 | 0 |
| 18.81 | 0.96 | 0.04 | 0 | 0 | 0 | 0 |
| 20.07 | 0 | 0.19 | 0 | 0 | 0.81 | 0 |
| 20.95 | 0.32 | 0.04 | 0.04 | 0.32 | 0.29 | 0 |
| 21.1 | 0.89 | 0.11 | 0 | 0 | 0 | 0 |
| 21.92 | 0 | 0.07 | 0 | 0 | 0.93 | 0 |
| 21.97 | 0.99 | 0.01 | 0 | 0 | 0 | 0 |
| 22.08 | 0 | 0 | 0 | 0 | 1 | 0 |
| 22.7 | 0.33 | 0.04 | 0 | 0.33 | 0.30 | 0 |
| 23.02 | 0.99 | 0.01 | 0 | 0 | 0 | 0 |
| 23.73 | 0 | 0 | 0 | 0.52 | 0.48 | 0 |
| 24.08 | 0 | 0 | 0 | 1 | 0 | 0 |
| 24.08 | 1 | 0 | 0 | 0 | 0 | 0 |
| 28.53 | 0.32 | 0.04 | 0.04 | 0.32 | 0.29 | 0 |
| 29.3 | 0.96 | 0.04 | 0 | 0 | 0 | 0 |
| 32.63 | 0.33 | 0.04 | 0 | 0.33 | 0.30 | 0 |
| 34.22 | 0.99 | 0.01 | 0 | 0 | 0 | 0 |
| 34.4 | 0 | 0 | 0 | 0 | 1 | 0 |
| 35.86 | 0.99 | 0.01 | 0 | 0 | 0 | 0 |
| 37.5 | 0 | 0 | 0 | 1 | 0 | 0 |
| 37.5 | 1 | 0 | 0 | 0 | 0 | 0 |
| 41.73 | 0 | 0.03 | 0 | 0 | 0.97 | 0 |
| 43.7 | 0.32 | 0.04 | 0.04 | 0.32 | 0.29 | 0 |
| 43.84 | 0 | 0.07 | 0 | 0 | 0.93 | 0 |
| 52.5 | 0.33 | 0.04 | 0 | 0.33 | 0.30 | 0 |
| 58.87 | 0.32 | 0.04 | 0.04 | 0.32 | 0.29 | 0 |
| 72.37 | 0.33 | 0.04 | 0 | 0.33 | 0.30 | 0 |
| 83.46 | 0 | 0.03 | 0 | 0 | 0.97 | 0 |
| 89.2 | 0.32 | 0.04 | 0.04 | 0.32 | 0.29 | 0 |
| 112.1 | 0.33 | 0.04 | 0 | 0.33 | 0.30 | 0 |
| 200 | 0 | 0 | 0 | 0 | 0 | 1 |

**A.**

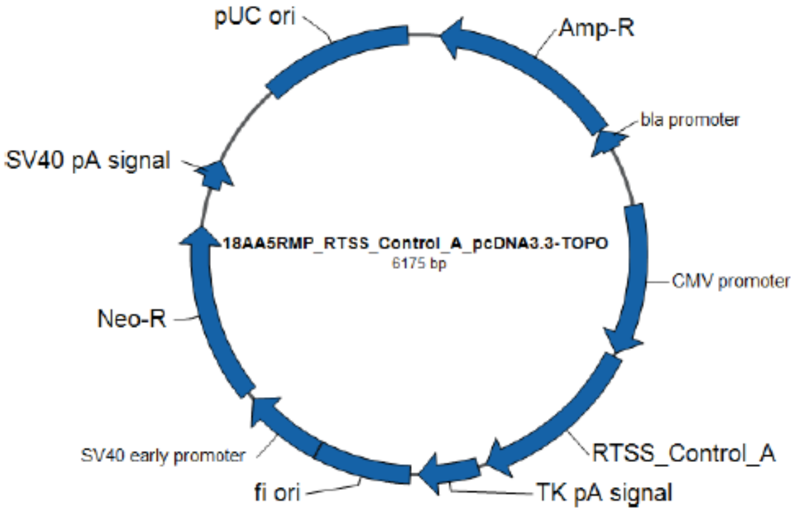


**B.**

GGATCCAAACTAAGATGTGTTCTTTATCTAA**TTAAGGAACAAGAAGGATAATACCA**TTATTAAT
CCTATTGAACTATCGATTACATTAAACACACTGGAACATTTTTCCATTTTACAAATTTTTTTT
CAATTCTTAATGCATAATCTAATTCGTCTTTAGGTTTATTAGCAGAGCCAGGCTTTATATCTCA
TTGAATACCATTTCCACAAGTTACACTACATGGGGACCGACTTATTGAAAGAGAATTTTGTATT
TTGTTTAAATATTCTTTTAGCTTGTTATCACTTGGTTCTTCGTTATTATTATTTTTTACAGCAC
TGTTGGCATTAGCATTTTCATCTA**CATTTCGGTTTGGGTCATTT**GGCATATTGTGACCTTGTCC
ATTACGGATCCTACAACTCCTCGAGGTAATCGTGGTAATTGTGGTCCTGC**TACTTTCCCTTGCC**
**CTTGTG**ATCCAGGTGATATCTTTGTTCCTACGGATCCGTTCCTTCCTGATGATCTTCCTGCTCT
TTCGACATCTGATTGGGATACTTCTGCACCTGGTCTTGCTGATTCTACTTTTTGTTCTGCTCCT
ATACCACTTCCTCTTCATTTTTGTTGTTGCTGTTGTTGGGGTTGTGTTTCTTGAGCTAAAGTTA
GATGTGTTGGTTGTTGTTTTGGTTTTTCAATTCTTGCTTGTGTCTATGTACGTGAACCATCGGA
TGATAATGA**TCCTGTAGCAGATTCATCTGTAGTG**GTTTCACTCTTTATTGTATTTTTACTCGAG

**Figure S1: Control plasmid map (A) and sequence (B).** The order of sequence in the plasmid is: Bam-25nt_linker-CSP_sequence-25nt_linker-Bam-8nt_linker-Xho-25nt_linker-SERA2_sequence-25nt_linker-Xho. Restriction sites are highlighted in yellow. Linker sequences are highlighted in grey. The primer sequences are marked in bold letters. Colored nucleotides denote naturally occurring SNPs observed in Neafsey *et al* ([1]; blue), the Pf3k database (www.malariagen.net/projects/pf3k; orange), or both (red). Underlined nucleotides mark regions of the plasmid sequence that differ from any previously observed natural sequences. Reference sequence nucleotide content was maintained within these regions, but the order of nucleotides were shuffled.

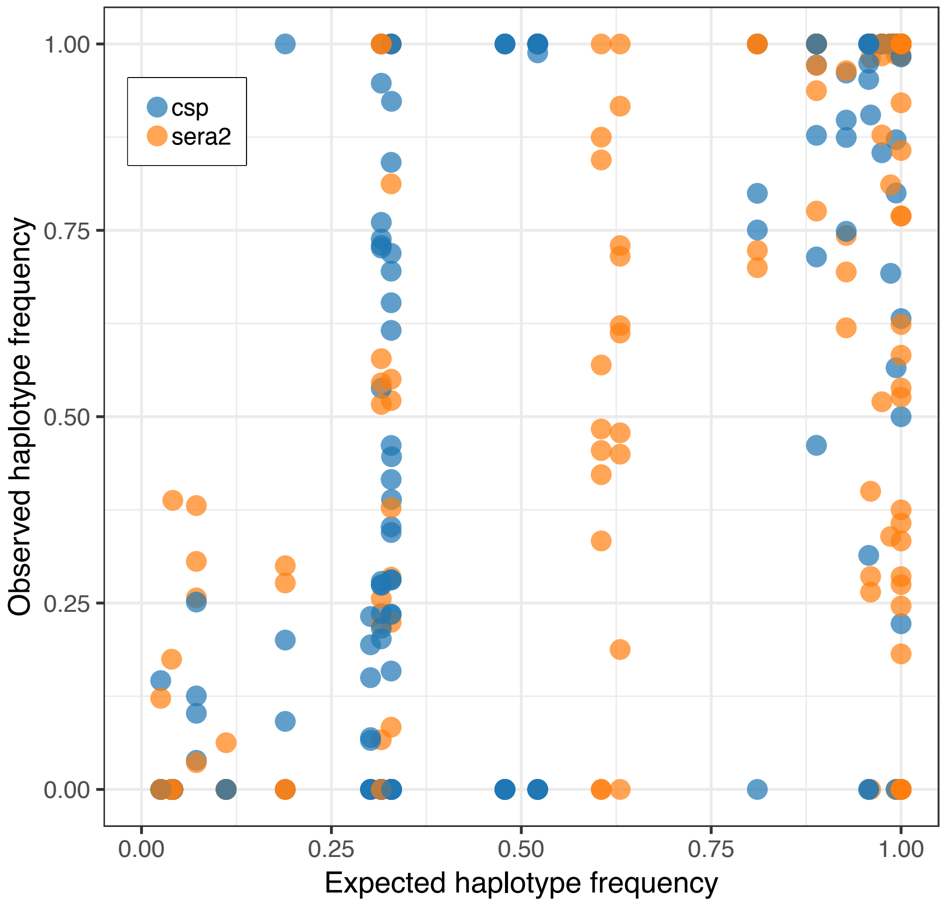


**Figure S2: Expected haplotype frequency versus observed haplotype frequency within samples.** Only samples with at least 100 reads are shown. Overall correlation (Pearson's r) is 0.82.

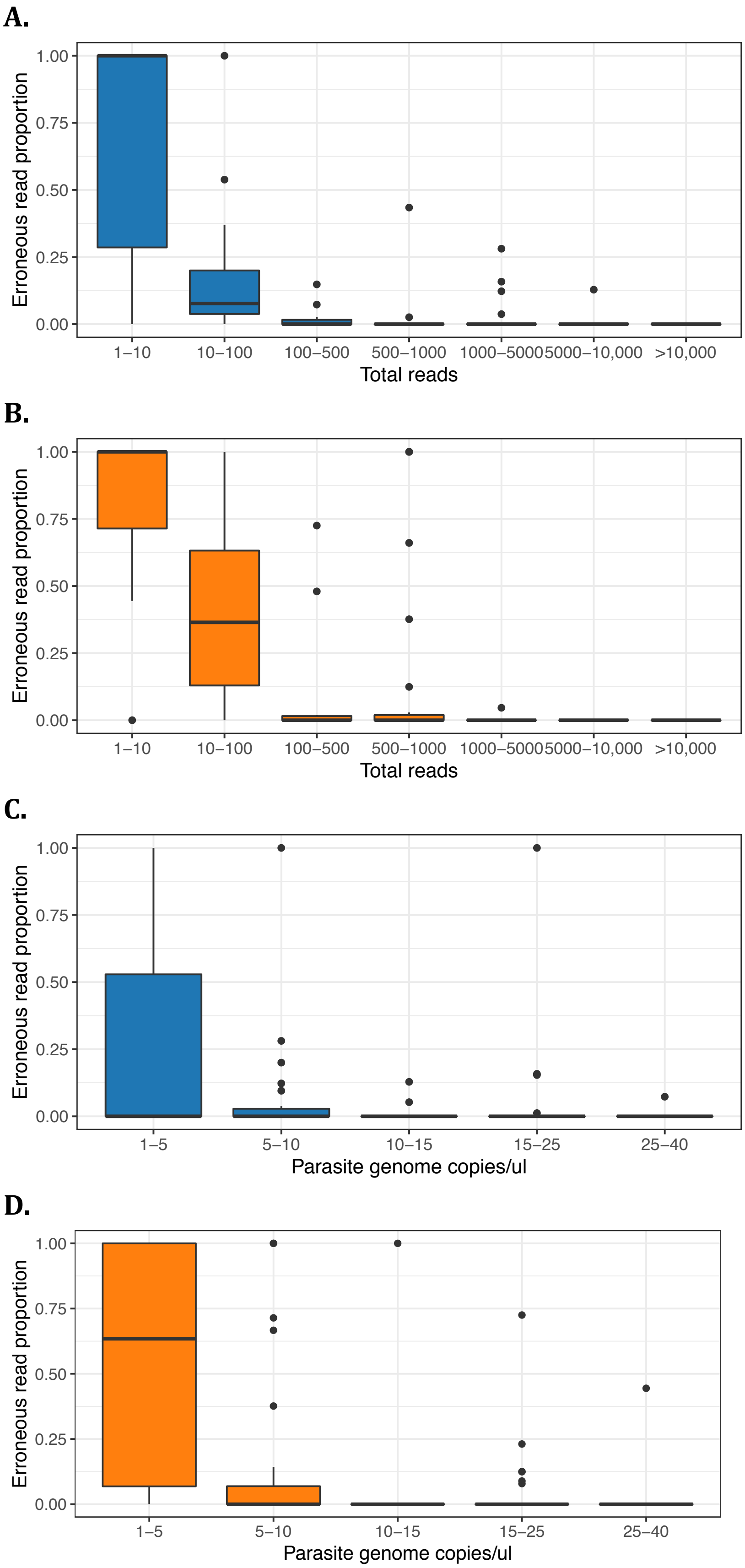


**Figure S3: Parasite DNA concentration and sample read depth affect the proportion of reads that are erroneous.** Samples with low read depth (A,B) and low parasite DNA concentrations (C,D) contain a higher proportion of reads that support erroneous haplotypes. Results are similar for the *CSP* (blue; A,C) and *SERA2* (orange; B,D) amplicons.

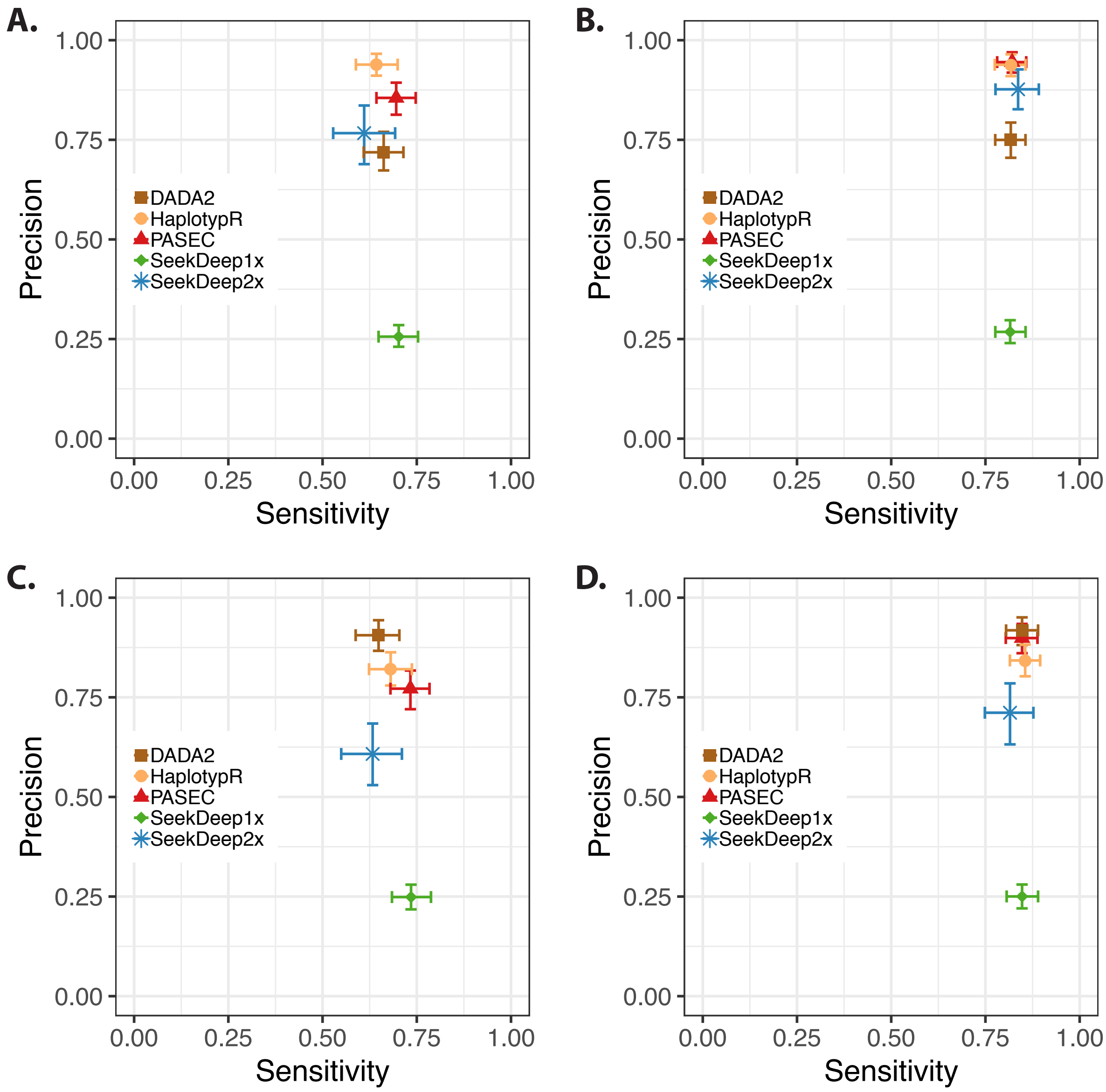


**Figure S4: Sensitivity and precision of the five analysis pipelines on a per amplicon basis.** Sensitivity and precision are plotted for *CSP* (A, B) and *SERA2* (C, D) The calculations for plots A. and C. were made using all samples, while plots B. and D. used only samples with at least 100 reads. 95% confidence intervals were calculated with 1000 bootstrap replicates.

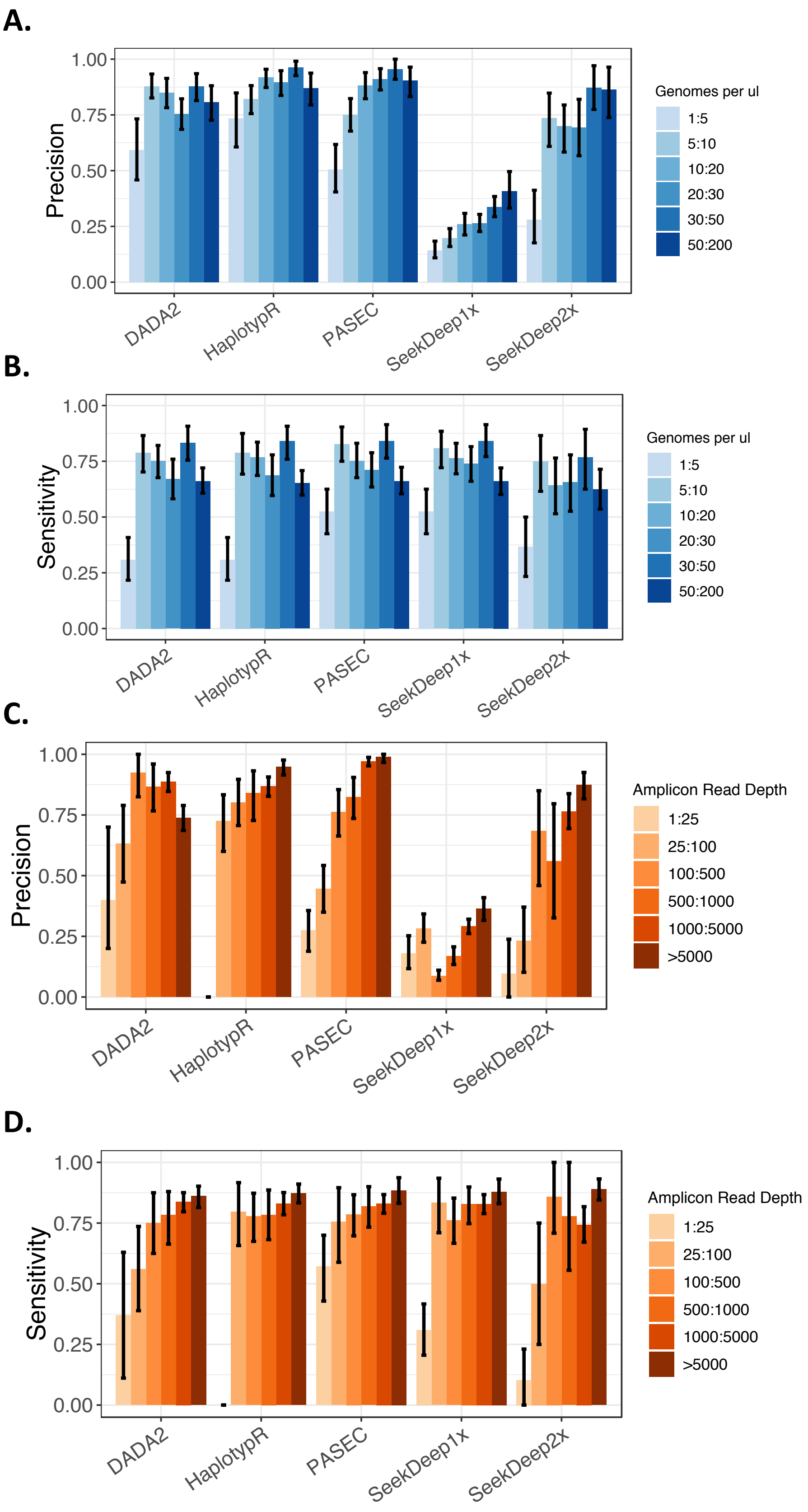


**Figure S5: Precision and Sensitivity are reduced at low genome copy number and low read depths.** HaplotypR best practices filter out all samples with fewer than 25 reads, so no data were available for this bin.

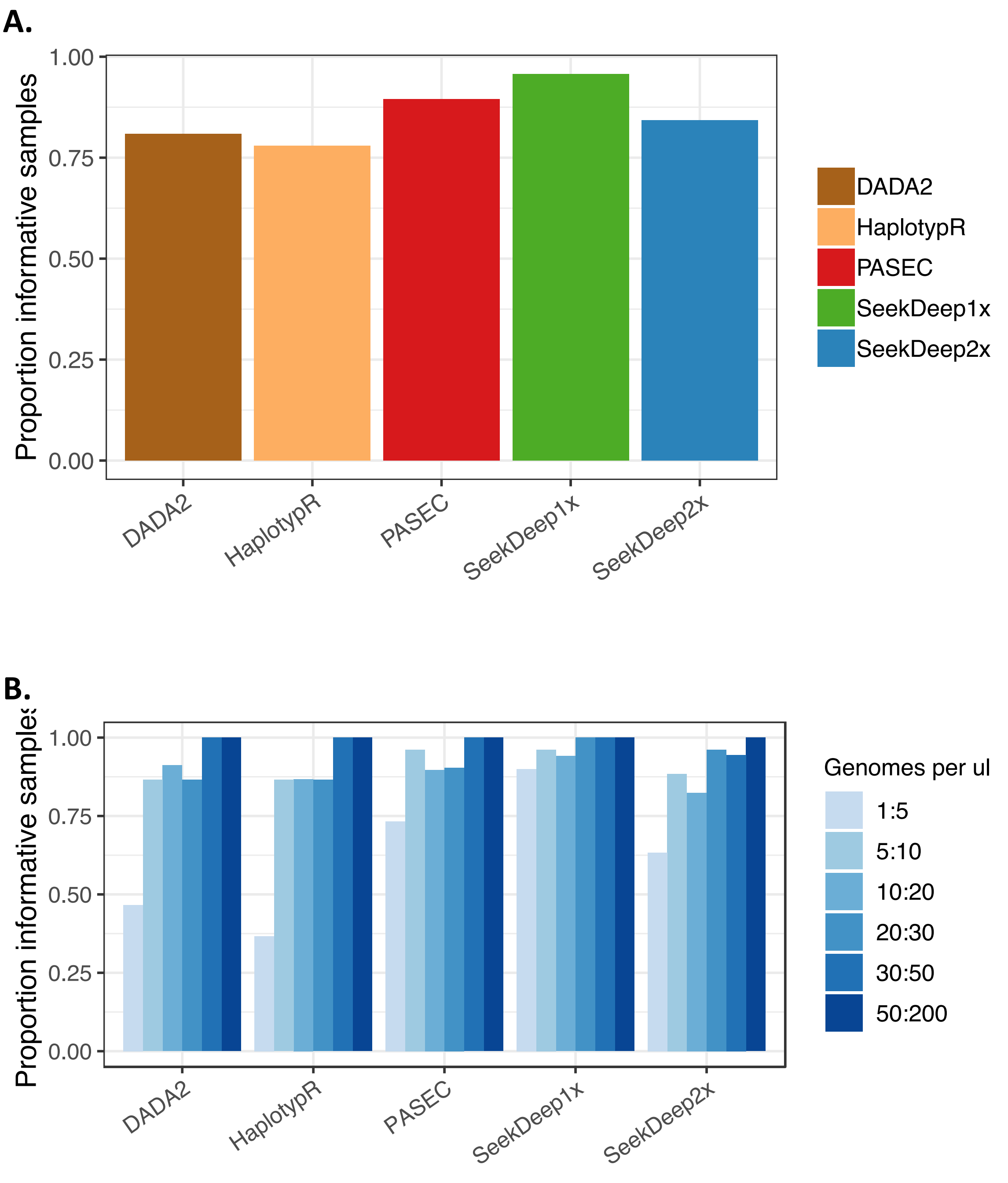


**Figure S6: Proportion of samples in which haplotypes were identified by each tool.** (A) Tools varied in their overall rate of sample success (the proportion of samples in which at least one haplotype was resolved). (B) Sample success was lowest for samples with low *Plasmodium* genome concentrations.

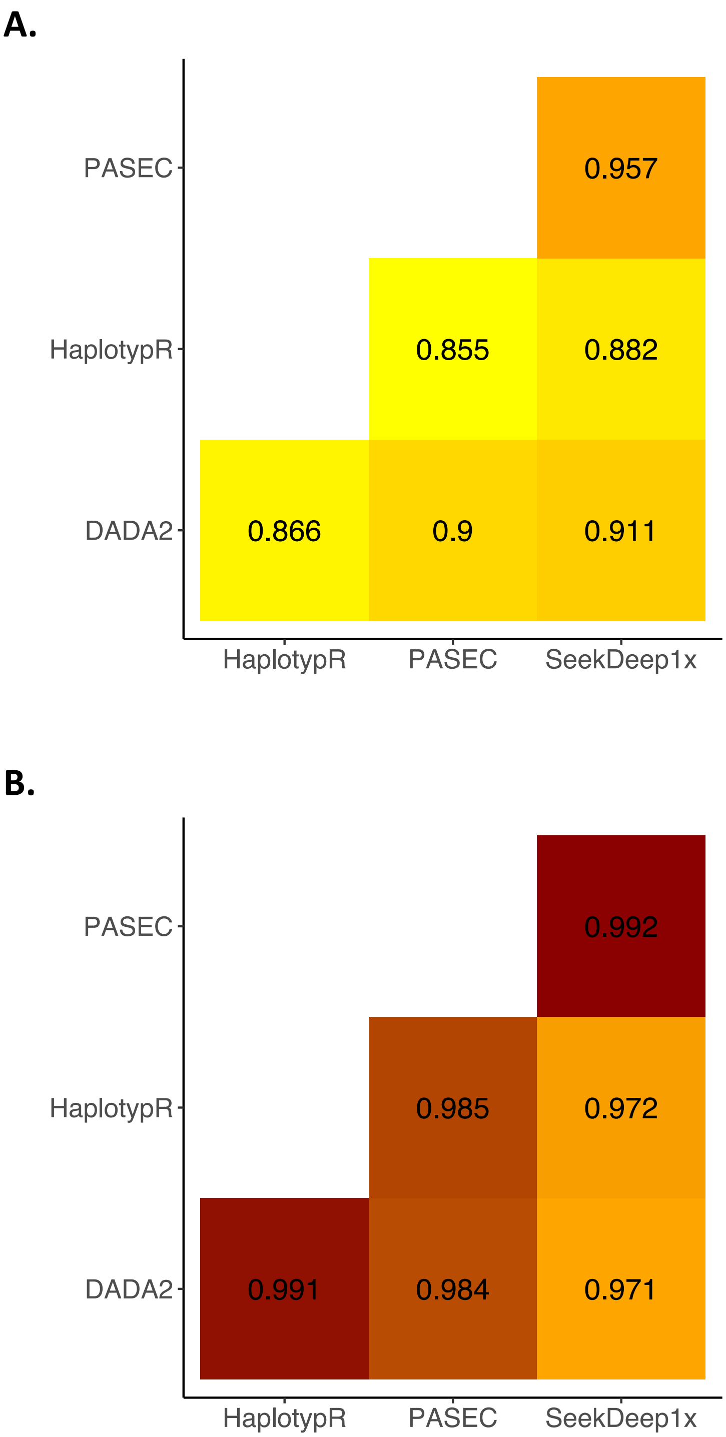


**Figure S7. Correlation between tool estimates of the major haplotype frequency within mock infections.** Pearson's r estimates are given for (A) all samples and (B) samples with at least 100 reads.

**A.**

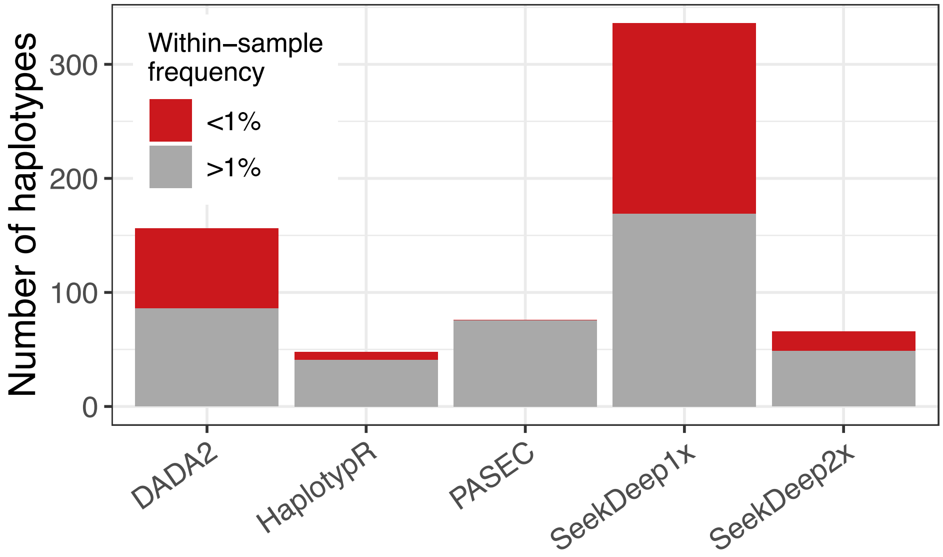


**B.**

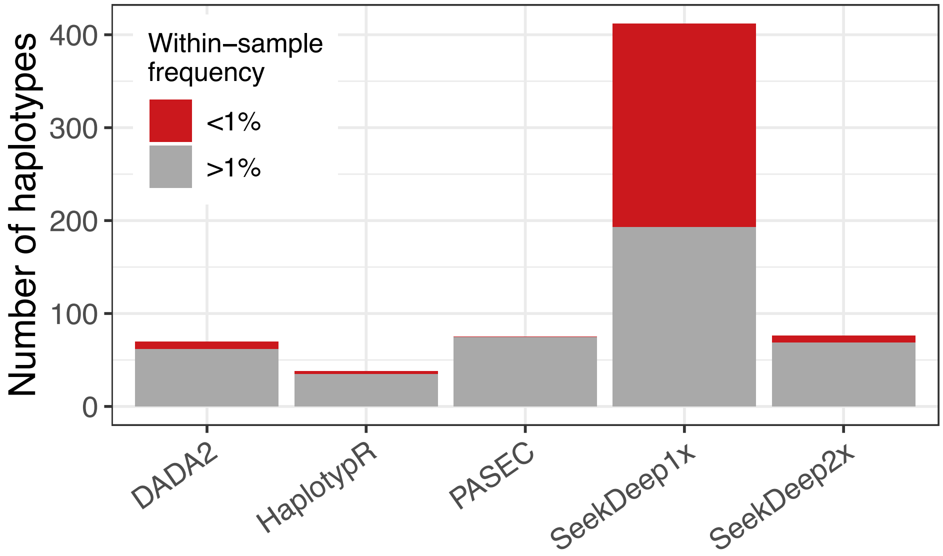


**Figure S8: Number of unique haplotypes identified by each tool in a set of 190 patient samples from sub-Saharan Africa.** Counts are shown for (A) *CSP* and (B) *SERA2*.

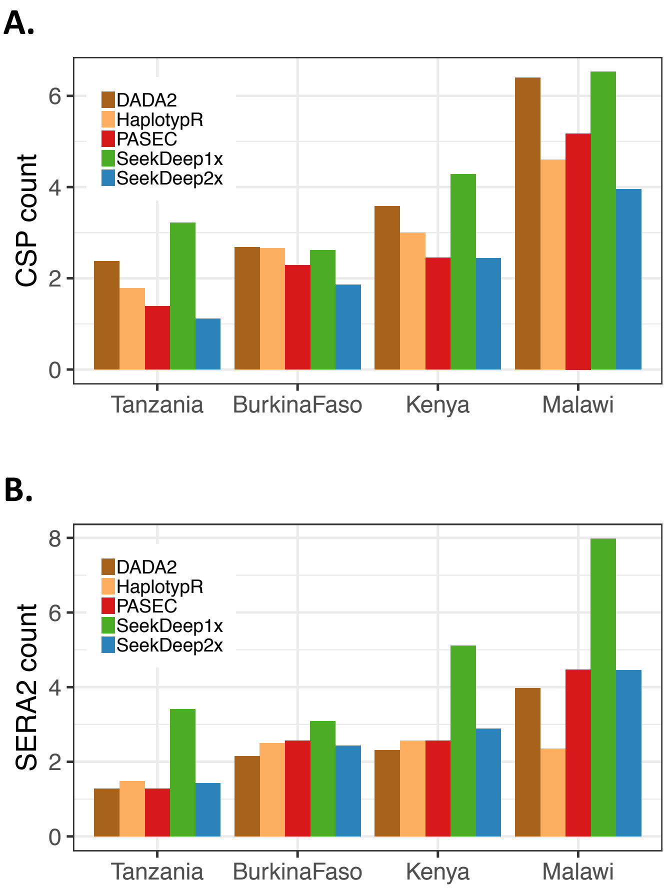


**Figure S9: Average number of (A) *CSP* and (B) *SERA2* haplotypes per sample calculated by the five pipelines.**

### References

1. Neafsey DE, Juraska M, Bedford T, Benkeser D, Valim C, Griggs A, et al. Genetic Diversity and Protective Efficacy of the RTS,S/AS01 Malaria Vaccine. N Engl J Med. 2015;373:2025–37. doi:10.1056/NEJMoa1505819.
