## Supplemental File 2 for "Detection of low-density *Plasmodium falciparum* infections using amplicon deep sequencing"

**Additional File 2: Supplementary Tables and Figures**

**Early *et al.*, Amplicon deep sequencing of low-density *Plasmodium falciparum* infections: an evaluation of analysis approaches**

### Parallel Amplicon Sequencing Error Correction (PASEC) workflow

The PASEC pipeline takes sample-demultiplexed BAM or FASTQ files, generates haplotypes for each BAM/ FASTQ in parallel and then pools those results into final tables after all threads have finished successfully (Figure A). It is implemented in Workflow Description Language (WDL) and utilizes the Cromwell execution engine, a workflow language and management system geared towards data-intensive scientific analysis pipelines (https://github.com/broadinstitute/cromwell).

For each individual BAM/FASTQ, paired end reads are first merged using FLASH [1] and then realigned to amplicon references using BWA-MEM [2]. After alignment, for each amplicon in parallel, reads are dereplicated, counted and optionally modified to produce a resultant haplotype. The user has several options for modifying the resulting haplotypes for customized error filtration. If the user wants to ignore indels, an 'ignore_indels' flag can be set to true in the configuration file. When enabled, deletion positions are padded with an 'X' symbol and all insertions are ignored. In addition, if the amplicons are sufficiently well-understood to identify homopolymeric regions known to be problematic for sequencing, the tool gives the option of passing a bed file specifying genomic positions to be masked.

After haplotypes are generated, the haplotype set is filtered according to a set of pre-specified thresholds. All haplotypes with length, coverage and intra-sample frequency under these thresholds are filtered from the analysis. After filtering, the haplotypes are sorted by coverage and clustered as follows. Starting with the highest coverage haplotype, each haplotype $H_i$ is pairwise compared to every other haplotype with less coverage $H_{j>i}$. Haplotype $H_j$ is clustered to $H_i$ if it is within some pre-specified edit and coverage-ratio distance of $H_i$, where edit distance is defined as $d = Levenshtein(H_i, H_j)$ and coverage-ratio as $r = \frac{coverage(H_j)}{coverage(H_i)}$, within a few restrictions. Haplotypes are allowed to belong to at most one cluster. If a haplotype is within the thresholds for multiple clusters, it will be clustered to the closest one (by Euclidean distance in edit distance and coverage ratio space) with highest coverage. Also, optionally, all haplotypes that have other haplotypes clustered to it are not eligible for clustering themselves. This effectively forces a maximum distance of twice the specified thresholds between any two haplotypes within a cluster.

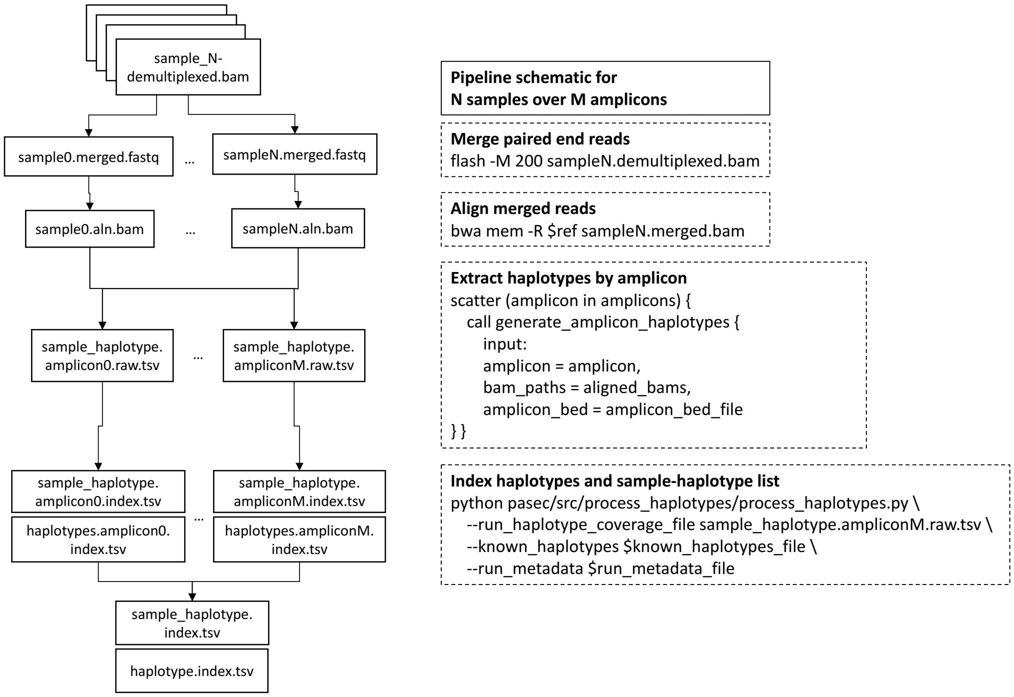


**Figure A. PASEC workflow**
