## Supplemental File 3 for "Detection of low-density *Plasmodium falciparum* infections using amplicon deep sequencing": supp_file_1-dada2.R

```
#
# amplicon seq validation - dada2 benchmark 
#
    
### imports
library(ggplot2)
library(reshape2)
library(optparse)
library(dada2); 

### useful fcns 
num_unique = function(x) sum(getUniques(x))

### prelims
# get paths from cmd-line args
option_list = list(make_option(c("-d", "--data_dir"), type="character", default=NULL, 
                               help="data directory path. required", metavar="character"),
                   make_option(c("-o", "--output_dir"), type="character", default=NULL, 
                               help="directory for output data. required", metavar="character"), 
                   make_option(c("-p", "--plot_dir"), type="character", default=NULL, 
                               help="directory for output plots. default is ${output_dir}/plots/", 
                               metavar="character")); 
opt_parser = OptionParser(option_list=option_list);
opt = parse_args(opt_parser);
if (is.null(opt$data_dir)){ 
    print_help(opt_parser)
    stop("--data_dir option required.", call.=FALSE)
}
if (is.null(opt$output_dir)) { 
    print_help(opt_parser)  
    stop("--output_dir option required.", call.=FALSE)
}
if (is.null(opt$plot_dir)) { 
    opt$plot_dir = file.path(opt$output_dir, "plots") 
}
# set read-types/ amplicons
read_types = c("R1", "R2")
amplicons = c("csp", "sera2")
# get list of fastqs
fastqs = list()
for (amplicon in amplicons) { 
    fastqs[[amplicon]] = list()
    for (read_type in read_types) {
        fastqs[[amplicon]][[read_type]] = sort(list.files(opt$data_dir, 
                                                          pattern=paste(amplicon, read_type, "fastq", sep='.'), 
                                                          full.names=TRUE))
    }
}
# set sample names
sample.names = unique(sapply(strsplit(basename(fastqs$csp$R1), '\\.'), '[', 1))

### filtering/ errors/ haplotypes 
## params
# truncation lengths for R1/ R2, respectively
# this parameter significantly effects filtering/ merging metrics downstream
truncation_lens = c(220, 200)

## data structs 
# filtering
filtering_stats = list()
filtered_fastqs = list()
# error 
error_profiles = list()
# haplotypes
haplotypes = list()
true_haplotypes = list()
merged_true = list()
haplotype_matrices = list()
haplotype_matrices.bimera_filtered = list()
haplotype_matrices_final = list()
# processing synopsis 
processing_stats = list()
processing_stats_percent = list()
processing_stats_melted = list()
processing_steps = c("input", "filtered", "clustered", "merged", "bimera_filtered")

## for each amplicon 
for (amplicon in amplicons) {
    ## plot quality profiles for forward/ reverse (R1/ R2) reads
    for (read_type in read_types) {
        for (i in 1:length(fastqs[[amplicon]][[read_type]])) { 
            sample = strsplit(basename(fastqs[[amplicon]][[read_type]][i]), '\\.')[[1]][1]
            g = plotQualityProfile(fastqs[[amplicon]][[read_type]][i]) + ggtitle(paste(sample, amplicon, read_type)) + 
                theme(plot.title = element_text(hjust = 0.5))
            ggsave(paste(paste0(opt$plot_dir, "/read_quality_profile"), sample, amplicon, read_type, "png", sep="."), plot=g)
        }
    }
    ## filter and trim 
    filtered_fastqs[[amplicon]] = list()
    filtered_fastqs[[amplicon]]$R1 = file.path(opt$data_dir, paste0(sample.names, '.', 
                                                                    amplicon, ".R1.filtered.fastq"))
    filtered_fastqs[[amplicon]]$R2 = file.path(opt$data_dir, paste0(sample.names, '.', 
                                                                    amplicon, ".R2.filtered.fastq"))
    filtering_stats[[amplicon]] = filterAndTrim(fastqs[[amplicon]]$R1, filtered_fastqs[[amplicon]]$R1,
                                                fastqs[[amplicon]]$R2, filtered_fastqs[[amplicon]]$R2, 
                                                truncLen=truncation_lens, maxN=0, rm.phix=TRUE, multithread=TRUE)
    # get those solexas which have zero reads after filtering
    zero_read_files = names(which(filtering_stats[[amplicon]][,'reads.out'] == 0))
    # remove those from the filtered_fastqs lists
    for (zero_read_file in zero_read_files) { 
        for (read_type in read_types) { 
            filtered_fastqs[[amplicon]][[read_type]] = names(which(sapply(filtered_fastqs[[amplicon]][[read_type]], 
                                                                          function(x) !grepl(strsplit(zero_read_file, '\\.')[[1]][1], x))))
        }
    }
    ## compute error and true haplotypes
    error_profiles[[amplicon]] = list()
    haplotypes[[amplicon]] = list()
    true_haplotypes[[amplicon]] = list()
    for (read_type in read_types) {
        # compute error profiles for R1/ R2, and plot
        error_profiles[[amplicon]][[read_type]] = learnErrors(filtered_fastqs[[amplicon]][[read_type]], 
                                                              multithread=TRUE)
        g = plotErrors(error_profiles[[amplicon]][[read_type]], nominalQ=TRUE) + 
            ggtitle(paste(amplicon, read_type)) + theme(plot.title = element_text(hjust = 0.5))
        ggsave(paste(paste0(opt$plot_dir, "/error_profiles"), amplicon, read_type, "png", sep="."), plot=g)
        # then haplotypes (i.e. unique reads w/ coverage)
        haplotypes[[amplicon]][[read_type]] = derepFastq(filtered_fastqs[[amplicon]][[read_type]])
        names(haplotypes[[amplicon]][[read_type]]) = sample.names
        # then infer true haplotypes, using denoising-clustering (i.e. dada)
        true_haplotypes[[amplicon]][[read_type]] = dada(haplotypes[[amplicon]][[read_type]],
                                                        err=error_profiles[[amplicon]][[read_type]],
                                                        multithread=TRUE)
    }
    ## merge R1/ R2 haplotypes 
    merged_true[[amplicon]] = mergePairs(true_haplotypes[[amplicon]]$R1,
                                         haplotypes[[amplicon]]$R1,
                                         true_haplotypes[[amplicon]]$R2,
                                         haplotypes[[amplicon]]$R2)
    ## init haplotype matrices, and filter any chimeras
    haplotype_matrices[[amplicon]] = makeSequenceTable(merged_true[[amplicon]])
    haplotype_matrices.bimera_filtered[[amplicon]] = removeBimeraDenovo(haplotype_matrices[[amplicon]],
                                                                        method="consensus", multithread=TRUE)
    ## save final haplotypes
    haplotype_matrices_final[[amplicon]] = melt(haplotype_matrices.bimera_filtered[[amplicon]])  
    colnames(haplotype_matrices_final[[amplicon]]) = c("solexa", "haplotype", "coverage")
    # filter rows w/ coverage == 0
    haplotype_matrices_final[[amplicon]] = 
        as.data.frame(haplotype_matrices_final[[amplicon]][haplotype_matrices_final[[amplicon]]$coverage > 0,])
    # sort by solexa_id 
    haplotype_matrices_final[[amplicon]] = 
        haplotype_matrices_final[[amplicon]][order(haplotype_matrices_final[[amplicon]]$solexa),]
    # save
    write.table(haplotype_matrices_final[[amplicon]], 
                file=paste(opt$output_dir, "/haplotypes", amplicon, "tsv", sep="."), sep="\t", 
                row.names=FALSE, quote=FALSE)
    ## plot synopsis of processing/ filtering steps
    # get raw numbers
    processing_stats[[amplicon]] = cbind(filtering_stats[[amplicon]], 
                                         sapply(true_haplotypes[[amplicon]]$R1, num_unique), 
                                         sapply(merged_true[[amplicon]], num_unique),
                                         rowSums(haplotype_matrices.bimera_filtered[[amplicon]]))
    colnames(processing_stats[[amplicon]]) = processing_steps
    rownames(processing_stats[[amplicon]]) = sample.names
    # convert to percentages 
    processing_stats_percent[[amplicon]] = t(apply(processing_stats[[amplicon]], 1, 
                                                   function(row) sapply(row, function(x) x / row[1])))
    colnames(processing_stats_percent[[amplicon]]) = processing_steps
    # plot 
    processing_stats_melted[[amplicon]] = melt(processing_stats_percent[[amplicon]])
    colnames(processing_stats_melted[[amplicon]]) = c("solexa", "processing_step", "percent_unfiltered")
    g = ggplot(data=processing_stats_melted[[amplicon]], aes(x=processing_step, y=percent_unfiltered, group=solexa)) + 
              geom_line(aes(color=solexa)) + ylim(0.0, 1.0) + theme_classic() + 
              ggtitle(amplicon) + theme(plot.title = element_text(hjust = 0.5))
    ggsave(paste0(opt$plot_dir, "/", amplicon, ".processing_steps.png"), plot=g)
}
```
