## Supplemental File 3 for "Detection of low-density *Plasmodium falciparum* infections using amplicon deep sequencing": supp_file_2A-haplotypR.sh

```
#!/usr/bin/env bash 
##
##  run_haplotypR.sh 
## 
##  calls haplotypR from command-line w/ default (clustering/filtering) settings 
##
##  where ~/Tools/HaplotypR/ is a local copy of https://github.com/tmfarrell/HaplotypR.git
## 

Rscript ~/Tools/HaplotypR/R/run_haplotypr.R \
    --output_dir=$1 --amplicons_file=$2 --samples_dir=$3 \
    --return_full_haplotypes --trim_reads --verbose
```
