## Supplemental File 3 for "Detection of low-density *Plasmodium falciparum* infections using amplicon deep sequencing": supp_file_3A-seekdeep.sh

```
#!/usr/bin/env bash 
set -e

data_dir=$1
output_dir=$2
sample_metadata=$3
primer_set_file=$4
group_metadata=$5
parameters_file=$6

if [ -d "$output_dir" ]; then 
    echo -e "\nAre you sure you want to remove \"${output_dir}\" before continuing? [y/n]:"
    echo "(The results of SeekDeep cannot be saved in a directory that already exists.)"
    read answer
    if [[ "$answer" == "y" ]]; then 
	rm -r $output_dir 
    else
	echo "Ok, choose another output_dir."
	exit
    fi
fi

# add SeekDeep to path 
export PATH=/Users/tfarrell/Tools/SeekDeep/bin/:$PATH

echo "SeekDeep: setting up analysis..."
SeekDeep setupTarAmpAnalysis --technology Illumina \
                             --inputDir $data_dir \
                             --outDir $output_dir \
                             --samples $sample_metadata \
                             --idFile $primer_set_file \
                             --groupMeta $group_metadata \
                             --extraExtractorCmds="--verbose" \
                             --extraQlusterCmds="--par ${parameters_file} --verbose" \
                             --extraProcessClusterCmds="--verbose"

cd $output_dir

echo "SeekDeep: extracting data..."
source extractorCmds.txt

echo "SeekDeep: qlustering haplotypes..."
source qlusterCmds.txt

echo "SeekDeep: process clusters..."
source processClusterCmds.txt
```
